# R-SPONDIN2^+^ Mesenchymal Cells Form the Bud Tip Progenitor Niche During Human Lung Development

**DOI:** 10.1101/2021.04.05.438484

**Authors:** Renee F.C. Hein, Joshua H. Wu, Yu-Hwai Tsai, Angeline Wu, Alyssa J. Miller, Emily M. Holloway, Tristan Frum, Ansley S. Conchola, Emmanuelle Szenker-Ravi, Bruno Reversade, Kelley S. Yan, Calvin J. Kuo, Jason R. Spence

## Abstract

Mammalian respiratory system development is regulated by complex reciprocal signaling events that take place between epithelial cells and the surrounding mesenchymal cells; however, mesenchymal heterogeneity and function in the developing human lung is poorly understood. We interrogated single cell RNA sequencing data from multiple human lung specimens and identified a mesenchymal cell population present during development that is highly enriched for expression of the WNT agonist *R-SPONDIN2* (*RSPO2*), and we found that adjacent epithelial bud tip progenitors are enriched for the RSPO2 receptor *LGR5*. By carrying out functional experiments using organoid models, lung explant cultures, and FACS-isolated *RSPO2*^+^ mesenchyme, we show that RSPO2 is a critical niche cue that potentiates WNT signaling in human lung progenitors to maintain their multipotency.

## INTRODUCTION

Lung development begins at approximately four weeks post conception in humans as the lung buds develop from the ventral anterior foregut endoderm. Soon after, the buds begin branching morphogenesis, leading to a network of epithelial tubes that include the trachea and series of bronchi that become progressively smaller, terminating at the gas-exchanging alveoli (Miller and Spence, 2017; Conway *et al*., 2020). During branching morphogenesis, the tip of each budding branch possesses a population of transient, epithelial progenitor cells called ‘bud tip’ progenitors. *In vivo* lineage tracing in animal models has shown that bud tip progenitors give rise to all cell types in the lung epithelium, including those that line the airways and the alveoli (Rawlins *et al*., 2009; Yang *et al*., 2018). More recently, a population of bud tip progenitors in the human lung has been identified, and functional experiments have shown that they have the ability to generate a broad spectrum of lung epithelial cell types (Nikolić *et al*., 2017; Miller *et al*., 2018, 2020). In mice, a specialized niche that supports bud tip progenitors is made up of surrounding mesenchyme that provides physical and biochemical support and determines whether bud tip progenitors will self-renew or differentiate into different epithelial cell types (Shu *et al*., 2002; Weaver, Batts and Hogan, 2003; Alejandre-Alcázar *et al*., 2007; Li *et al*., 2008; Rajagopal *et al*., 2008; Tsao *et al*., 2008; Goss *et al*., 2009; Morrisey and Hogan, 2010; McCulley, Wienhold and Sun, 2015; Zepp and Morrisey, 2019; Riccetti *et al*., 2020). Genetic gain- and loss-of-function studies have identified many of the signaling pathways important for progenitor maintenance and for determining cell-fate choices during differentiation (Morrisey and Hogan, 2010). Drawing from these studies, a minimal set of essential signaling cues required to maintain isolated human bud tip progenitor cells in long-term *in vitro* culture has recently been described (Nikolić *et al*., 2017; Miller *et al*., 2018); however, the specific mesenchymal cells and signaling components that make up the *in vivo* bud tip progenitor niche are unclear. Moreover, we have only begun to scratch the surface in identifying the mechanisms that are conserved between animal models and humans (Danopoulos, Shiosaki and Al Alam, 2019; Conway *et al*., 2020; Miller *et al*., 2020).

Here, we investigate mesenchymal cell populations in the developing human distal lung from 8 – 19 weeks post-conception, a time when the lung supports an actively branching bud tip progenitor population. Leveraging single cell RNA sequencing (scRNA-seq) data and unsupervised clustering analysis, we identified transcriptionally distinct mesenchymal cell populations in the distal lung domain during this time frame, including smooth muscle cells and three, non-smooth muscle mesenchymal cell clusters that are highly enriched for expression of the WNT agonist *R-SPONDIN 2* (*RSPO2*). It is notable that *Rspo2*^*-/-*^ mice have mild lung defects compared to lung aplasia that is seen in humans with *RSPO2* mutations (Bell *et al*., 2008; Szenker-Ravi *et al*., 2018). Indeed, mutations in human *RSPO2* are lethal (Szenker-Ravi *et al*., 2018); however, the specific role of RSPO2 in the developing human lung has not been interrogated.

Based on the severe phenotype linked with *RSPO2* mutations in humans as well as the large population of *RPSO2*^+^ cells identified in scRNA-seq data, we interrogated the spatial localization of *RPSO2*^+^ cells along with the functional role of RSPO2 using human tissue, explant culture systems, and organoids. By fluorescence *in situ* hybridization (FISH) and immunofluorescence (IF), we show that *RSPO2* is expressed in mesenchymal cells physically located adjacent to bud tip progenitors. On the other hand, SM22^+^ airway smooth muscle cells lack *RSPO2* expression and line the newly differentiating proximal (airway) epithelium in domains adjacent to bud tip progenitor cells. In addition, we found that the RSPO2 receptor *LGR5*, but not other *LGR* family members, is uniquely expressed in bud tip progenitors, as are other canonical WNT target genes such as *AXIN2*. Using *in vitro* lung explant cultures with functional inhibition experiments, we show that blocking endogenous RSPO signaling leads to a loss of the high WNT signaling environment in the bud tip domain and stochastic differentiation of bud tip progenitors into multiple proximal, but not distal, lung cell types. Lastly, we identified LIFR as a cell surface marker for *RSPO2*^+^ cells and show that FACS-isolated LIFR^HI^ cells support the ability of bud tip organoids to give rise to both proximal and distal epithelium while LIFR^-^ cells only support proximal differentiation in co-cultures. Collectively, this work identifies an RSPO2-producing niche cell in the human fetal distal lung mesenchyme and reveals a critical RSPO2*-* mediated WNT signaling axis that supports bud tip progenitor cell maintenance and multipotency throughout early lung development.

## RESULTS

### Single cell RNA sequencing identifies mesenchymal cell populations in the fetal distal lung

To interrogate the mesenchymal cells present in the developing human lung in the bud tip progenitor domain, we re-analyzed scRNA-seq data from the physically-isolated distal portion of 8 – 19 week post-conception lungs (n = 5 lungs) (Figure 1A) (Miller *et al*., 2020). We used principal component analysis (PCA) for dimensionality reduction, Louvain clustering, and UMAP for visualization (Wolf, Angerer and Theis, 2018; Becht *et al*., 2019). Expression analysis of canonical marker genes identified major cell classes within the data, including epithelial, mesenchymal, immune, and endothelial cell types (Figures S1A and S1B). In order to specifically interrogate mesenchymal cell types that may comprise the bud tip progenitor niche, we extracted and re-clustered the mesenchymal cell clusters (clusters 0, 1, and 2), which were defined by expression of *VIM, POSTN, DCN, TCF21, COL1A1*, and *COL3A1*. Vascular smooth muscle cells (cluster 4) were excluded from the re-clustering analysis because they were identified as a clear, independent population expressing *TAGLN, ACTA2*, and *PDGFRB* (Figures S1A and S1B).

**Figure 1.**
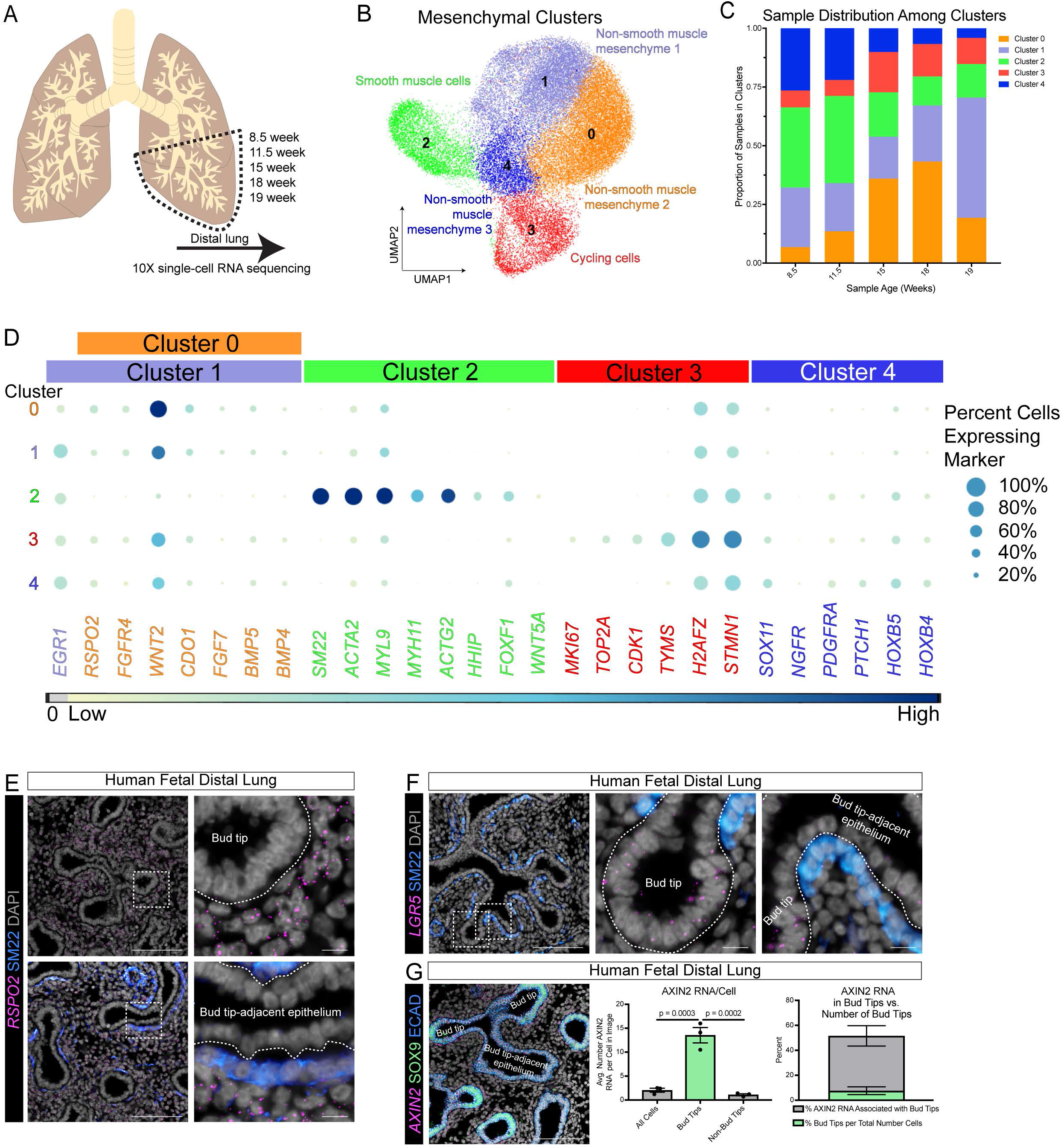
Identification of bud tip-associated mesenchymal populations and the close association of *RSPO2*^+^ mesenchymal cells with bud tip progenitors. (A) Schematic showing sample ages (days post-conception) and the general location of the distal lung where samples were taken from for single cell RNA sequencing. (B) Cluster plot of non-vascular smooth muscle mesenchymal cells (*VIM*^+^/*POSTN*^+^/*DCN*^+^*/TCF21*^*+*^/*COL1A2*^+^/*COL3A1*^+^/*SM22*^*+*^/*ACTA*^+^/*PDGFRB*^*-*^). Each dot represents a single cell and cells were computationally clustered based on transcriptional similarities. The plot is colored and numbered by cell-type identity of the cells composing each cluster. Cell-type labels for each cluster are based on expression of canonical (smooth muscle cells and cycling cells) or novel (non-smooth muscle mesenchyme) cell-type markers displayed in the dot plot in Figure 1D. We identified four distinct mesenchymal cell clusters and one cluster defined by markers of proliferation (cluster 3). Cluster 2 represents airway smooth muscle cells, and we defined the other three clusters (clusters 0, 1 and 4) as non-smooth muscle mesenchyme. (C) Stacked bar graph displaying the proportion of cells from each sample in each cluster of the cluster plot in Figure 1B. Cluster 0 is dominated by the 15- and 18-week samples with low contributions from the 8.5-. 11.5, and 19-week samples. Cluster 1 is dominated by the 19-week sample with smaller contributions from the remaining samples. Cluster 2 has large contributions from the 8.5- and 11.5-week samples with smaller contributions from the 15-, 18-, and 19-week samples. Cluster 3 has contributions from each sample, with higher contributions from the 15-, 18-, and 19-week samples. Cluster 4 is dominated by the 8.5- and 11.5-week samples with small contributions from the 15-, 18-, and 19-week samples. (D) Dot plot of genes enriched in each cluster of the UMAP plot in Figure 1B. The dot size represents the percentage of cells expressing the gene in the corresponding cluster, and the dot color indicates log-normalized and z-transformed expression level of the gene. Clusters are colored corresponding to the cluster plot in Figure 1B. (E) Fluorescence *in situ* hybridization of *RSPO2* and co-immunofluorescence for SM22 on 12-week human fetal distal lung tissue sections. DAPI is shown in gray. Scale bars represent 100µm or 10µm for insets. *RSPO2* is expressed broadly throughout the mesenchyme, and *RSPO2*^*+*^ cells sit adjacent to bud tips (top images). SM22^+^ cells line bud tip-adjacent epithelium and is negative for expression of *RSPO2* (bottom images). All epithelium is negative for *RSPO2* expression. See Figure S1 for *RSPO2* expression in the proximal lung and expression of other *RSPO* transcripts. (F) Fluorescence *in situ* hybridization of *LGR5* and co-immunofluorescence for SM22 on 17.5-week human fetal distal lung tissue sections. DAPI is shown in gray. Scale bars represent 100µm or 10µm for insets. *LGR5* expression is localized to bud tip progenitors and is largely absent from non-bud tip epithelium and mesenchyme in the distal lung. See Figure S1 for *LGR5* expression in the proximal lung and for expression of other *LGR* transcripts. A) The left-most image shows fluorescence *in situ* hybridization for *AXIN2* and co-immunofluorescence for bud tip marker SOX9 on 13.5-week human fetal distal lung tissue sections. DAPI is shown in gray. Scale bar represents 100µm. The graphs show that *AXIN2* expression is enriched in bud tip progenitors compared to other cells in the distal lung. The middle graph shows that bud tips have an average of 13.5 *AXIN2* RNA molecules per cell compared to an average of 1.1 *AXIN2* RNA molecules in non-bud tip cells (p = 0.0002, ordinary one-way ANOVA) and an average of 2.0 *AXIN2* molecules in all cells (p = 0.0003, ordinary one-way ANOVA) in a single-plane image of a 4µm tissue section. The right-most graph shows that even though bud tips only make up approximately 7.6% of cells in the distal lung, approximately 43.9% of *AXIN2* RNA molecules are associated with bud tip progenitor cells. Error bars show standard error of the mean. Each data point represents a separate image field from the same tissue specimen.

Re-clustering identified 5 transcriptionally distinct mesenchymal cell clusters (Figure 1B). Cluster 3 consisted of cells expressing proliferation genes, including *MKI67, TOP2A*, and others (Figures 1D). Differential expression analysis showed that cluster 2 contains cells expressing canonical markers of smooth muscle cells (e.g., *SM22, ACTA2, MYL9, MYH11, ACTG2*) and is enriched for expression of *HHIP, FOXF1*, and *WNT5A* (Figures 1D). Cells in clusters 0, 1, and 4 were identified as non-smooth muscle mesenchymal cells and had very similar gene expression profiles, expressing enriched levels of *RSPO2, FGFR4*, and *WNT2*, among others (Figures 1D). Although cells from these three clusters share the expression of many genes, there were subtle differences. For example, cells in cluster 4 also share enrichment for genes notable in the smooth muscle cell cluster compared to clusters 0 and 1, such as *SOX11, NGFR, PDGFRA, PTCH1, HOXB5*, and *HOXB4* while clusters 0 and 1 differ in expression of *EGR1* (Figures 1D). Of note, transcriptional differences between clusters 0, 1, and 4 may largely be based on gestational age, since we observed a nonequivalent distribution over time (Figure 1C), with the 8.5- and 11.5-week samples contributing most to clusters 2 and 4, the 15- and 18-week samples contributing most to cluster 0, and the 19-week sample contributing most to cluster 1 (Figure 1C). In addition, although *EGR1*^+^/*RSPO2*^*+*^ and *EGR1*^*-*^/*RSPO2*^*+*^ mesenchymal cells were found in the same tissue section by fluorescence *in situ* hybridization (FISH), expression of *EGR1* clearly increases over gestational age (Figure S1C). Of note, the three non-smooth muscle mesenchymal clusters express the WNT signaling molecules *WNT2* and *RSOP2* (Figures S1D). Given that the growth of human bud tip progenitor cells in culture requires exogenous WNT stimulation (Nikolić and Rawlins, 2017; Miller *et al*., 2018), we hypothesized these cells might make up an important part of the bud tip progenitor niche.

### *RSPO2*^*+*^ mesenchymal cells are localized adjacent to bud tip progenitor cells

In order to spatially profile *RSPO2*^+^ mesenchymal cells, we used FISH combined with IF on human fetal lung tissue sections spanning 8 – 19 weeks post-conception. *RSPO2*^*+*^ mesenchymal cells were co-visualized with airway smooth muscle cells (SM22^+^) because the scRNA-seq data showed that these populations are molecularly distinct. Co-FISH/IF for *RSPO2* and SM22 confirmed that these markers are expressed in different mesenchymal cell populations. *RSPO2*^*+*^ mesenchymal cells are located physically adjacent to bud tip progenitor cells while SM22^+^ airway smooth muscle cells line the more proximal, bud tip-adjacent epithelium (Figure 1E). Additional FISH data showed that *RSPO2*^*+*^ cells express *WNT2* and *FGFR4*, confirming scRNA-seq data and supporting previous reports revealing expression of *WNT2* and *FGFR4* in the distal lung near bud tip progenitors in mice and humans (Figure S1D) (Goss *et al*., 2009; Miller *et al*., 2012; Danopoulos *et al*., 2018; Danopoulos, Shiosaki and Al Alam, 2019; Yu *et al*., 2020). Also in agreement with scRNA-seq data, *PDGFRa* expression was found in both *RSPO2*^+^ cells and in SM22^+^ cells and appeared particularly enriched in SM22^+^ cells directly adjacent to bud tip regions (Figure S1E). In contrast to the distal lung domain, *RSPO2* expression is nearly absent from lung mesenchyme surrounding the bronchi and trachea and from all epithelium (Figure 1E and S1F). Based on scRNA-seq and FISH, the other *R-SPONDIN* transcripts were detected at much lower levels compared to *RSPO2* and not specifically localized near bud tip or differentiating epithelium (Figure S1G). The proximity and specificity of *RSPO2*^*+*^ mesenchyme to bud tip progenitors further suggested that RSPO2 comprises an important component of the bud tip progenitor niche.

### *LGR5* is expressed in bud tip progenitor cells

One mechanism by which R-SPONDIN proteins are known to amplify WNT signaling is by binding to LGR receptors and sequestering ubiquitin ligases that act on WNT receptor complexes, subsequently freeing the WNT receptor Frizzled from protein degradation (de Lau, Snel and Clevers, 2012; Niehrs, 2012; Chen *et al*., 2013; de Lau *et al*., 2014; Park *et al*., 2018; Raslan and Yoon, 2019). To determine which cells RSPO2^+^ mesenchymal cells may signal to, we used FISH to characterize the localization of *LGR* receptors within the fetal lung. *LGR5* (but not *LGR4* or *LGR6*) is highly specific to bud tip progenitor cells in the distal lung (Figure 1F). *LGR5* expression is largely excluded from mesenchymal cells and from differentiated epithelial cell types, except for a subset of basal cells in the proximal airways (Figure S1H). In contrast, *LGR4* is expressed broadly throughout the mesenchyme in the distal and proximal lung and is excluded from the epithelium while *LGR6* is expressed specifically in airway smooth muscle cells in the distal and proximal lungs (Figure S1H). The specific expression of *LGR5* in bud tip progenitors suggests that LGR5 may be a cognate receptor for *RSPO2* present in the bud tip progenitor niche.

### WNT target gene expression is enriched in bud tip progenitor cells

Given the expression pattern of *RSPO2* and *LGR5*, we predicted bud tip progenitors would display higher levels of WNT-mediated target gene expression compared to other cell types in the distal lung. Using *AXIN2* expression as a read-out for WNT signaling, quantification of FISH data revealed that *AXIN2* is enriched in bud tip progenitor cells compared to other cell types in the distal lung (Figure 1G). Using quantitative image analysis, we determined that in a single-plane image of a 4µm tissue section, bud tips had an average of 13.5 *AXIN2* RNA molecules per cell while the average number of *AXIN2* RNA molecules in all other cells was 1.1 (Figure 1G, middle panel). We also found that although bud tips only make up approximately 7.6% of the total number of cells in an image, they contain nearly 43.9% of the *AXIN2* RNA molecules in the image (Figure 1G, rightmost panel). This data shows that bud tip progenitors in the human fetal distal lung have enriched WNT target gene expression. Based on our collective data, this further supports our hypothesis that RSPO2 from the mesenchyme may act on bud tip progenitors via LGR5 to support a high WNT signaling domain to maintain bud tip progenitors.

### RSPO2-mediated WNT signaling in bud tips is required for proximal-distal patterning

To test the necessity of RSPO2-mediated WNT signaling for bud tip progenitor maintenance, we used an adenovirus (ad) expressing the soluble ectodomain of LGR5 (hereafter termed LGR5 ECD), which was previously shown to bind and neutralize RSPO2 leading to reduced levels of WNT signaling (Yan *et al*., 2017). We infected human fetal lung explants placed in an air-liquid-interface culture system with the LGR5 ECD ad or a control adenovirus encoding murine immunoglobulin IgG2a (Yan *et al*., 2017) every 2 days for 4 days. We confirmed successful infection of the virus via antibody staining against murine IgG2a_Fc for the control and against FLAG for the LGR5 ECD (Figure S2A). The explants continued to grow over this culture period (Figure 2B) with no significant differences in KI67 expression between the control and LGR5 ECD ad-infected explants (Figure S2B). In some cases, independent of viral infection, explants grew abnormally large, leading to cell death or necrosis towards the center of the explant; however, Cleaved Caspase 3 (CCASP3) staining was low or absent in most explants (Figure S2C). In addition, because both viruses appeared to infect the edge of the explants more strongly than the center (Figure S2A), we focused our analysis to the periphery of explants.

**Figure 2.**
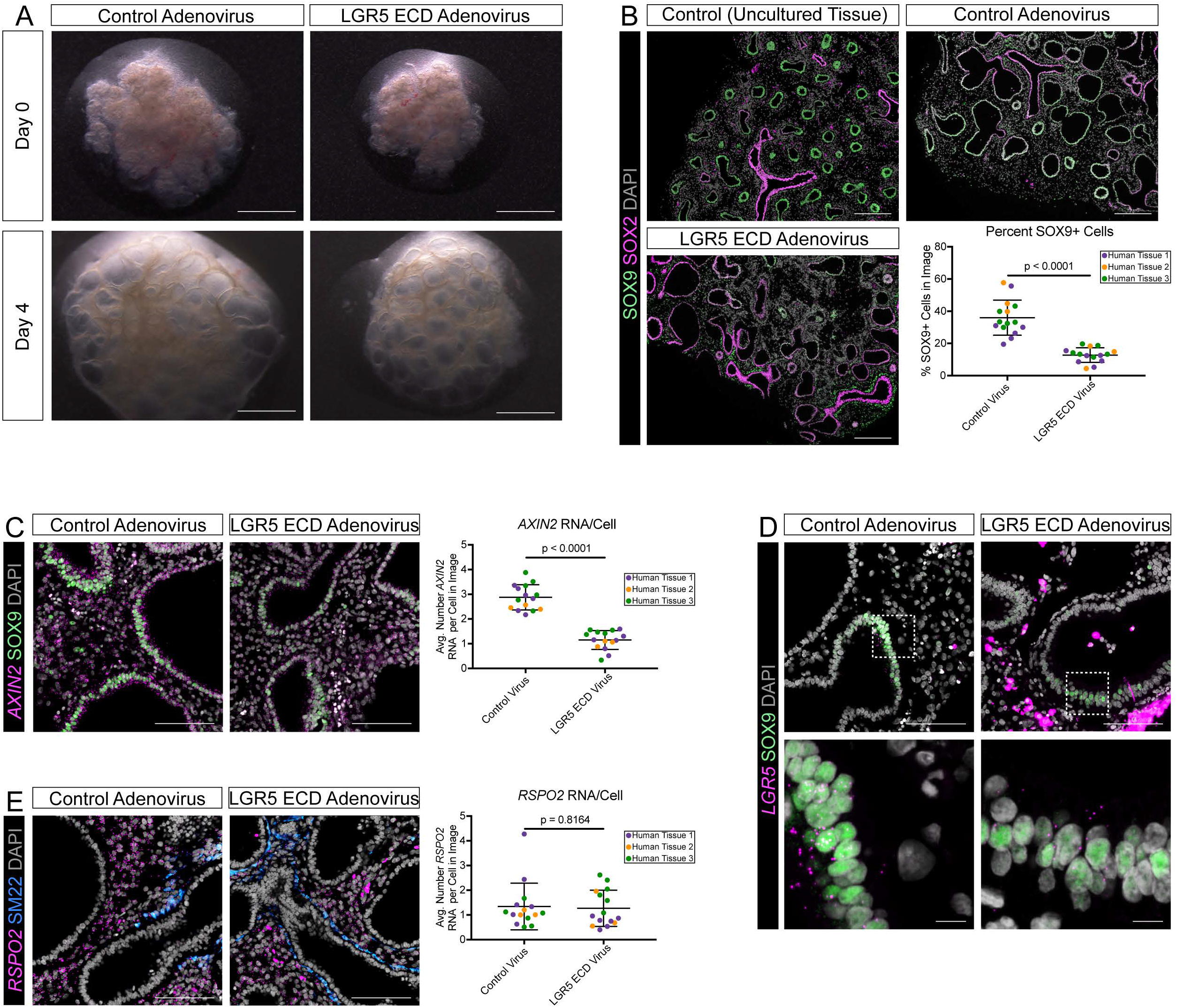
Inhibition of RSPO2-mediated WNT signaling in lung explants disrupts proximal-distal patterning. (A) Brightfield images of 11.5-week human fetal lung explants at the start of air-liquid interface culture and after 4 days of culture with 2 infections of either a control or LGR5 ECD adenovirus (ad). Scale bars represent 1mm. Both the control and the LGR5 ECD ad-infected explants grew over the 4-day culture period. In both conditions, mesenchymal cells were retained and the epithelium expanded. (B) Immunofluorescence for bud tip/distal marker SOX9 and proximal lung epithelial marker SOX2 on sections from uncultured human fetal lung tissue, control ad-infected explants, and LGR5 ECD ad-infected explants. DAPI is shown in gray. Scale bars represent 200µm. In the uncultured tissue, SOX9 is expressed highly in bud tip progenitor cells while SOX2 is expressed highly in proximal airway cells and lower in bud tip progenitor cells as previously described (Nikolić *et al*., 2017; Miller *et al*., 2018). The control-ad infected explants retained SOX9 expression in the bud tip regions with proper SOX9/SOX2 proximal-distal patterning while the bud tip regions in the LGR5 ECD ad-infected explants began to lose SOX9 expression. Quantification of the SOX9 stain is shown in the bottom right. At the end of the 4-day culture period, SOX9^*+*^ cells in the control ad-infected explants comprised approximately 36.0% of cells while they only comprised 12.8% in LGR5 ECD ad-infected explants (p < 0.0001, Welch’s t test). This quantification was performed in three unique biological samples with one to three technical replicates and a minimum of three image fields for each sample. (C) Fluorescence *in situ* hybridization and quantification of *AXIN2* and co-immunofluorescence for bud tip/distal marker SOX9 on control ad-infected explants and LGR5 ECD ad-infected explants. DAPI is shown in gray. Scale bars represent 100µm. In control ad-infected explants, there was an average of 2.9 *AXIN2* molecules per cell in a single-plane image while only an average of 1.2 *AXIN2* molecules per cell in a single-plane image in LGR5 ECD ad-infected explants (p < 0.0001, Welch’s t test). This quantification was performed in three unique biological samples with one to three technical replicates and a minimum of three image fields for each sample. (D) Fluorescence *in situ* hybridization (FISH) of *LGR5* and co-immunofluorescence for bud tip/distal marker SOX9 on sections from control ad-infected explants and LGR5 ECD ad-infected explants. DAPI is shown in gray. Scale bars represent 100µm or 10µm for insets. Endogenous *LGR5* expression was retained in both the control ad-infected explants and the LGR5 ECD ad-infected explants and appears restricted to bud tip progenitors. Viral *LGR5* was also detected by the FISH probe throughout the LGR5 ECD ad-infected explants (pink non-punctate stain). Expression of *LGR5* in bud tip progenitors appears decreased in LGR5 ECD ad-infected explants compared to control ad-infected explants; however, endogenous *LGR5* expression was quantified due to probe detection of viral *LGR5*. (E) Fluorescence *in situ* hybridization and quantification of *RSPO2* and co-immunofluorescence for smooth muscle marker SM22 on sections from control ad-infected explants and LGR5 ECD ad-infected explants. DAPI is shown in gray. Scale bars represent 100µm. The level of *RSPO2* expression in LGR5 ECD ad-infected explants is not significantly different than *RSPO2* expression in control ad-infected explants (p = 0.8164, Welch’s t test), and *RSPO2* expression remained absent from SM22^*+*^ smooth muscle cells in both conditions. This quantification was performed in three unique biological samples with one to three technical replicates and a minimum of three image fields for each sample.

Following 4 days of culture, the explants infected with the LGR5 ECD exhibited reduced staining for the bud tip marker SOX9 in the epithelium, with SOX9^+^ cells making up 36.0% of total cells in the control but only 12.8% of cells in the LGR5 ECD ad-infected explants (Figure 2B). In comparison to *in vivo*, uncultured lung tissue of a similar gestational age and previous reports (Abler *et al*., 2017; Miller *et al*., 2018), the control ad-infected explants maintained proper SOX2 and SOX9 epithelial patterning, with SOX9^HI^/SOX2^LOW^ bud tips and SOX9^-^/SOX2^HI^ proximal epithelium (Figure 2B, top). Much of the epithelium in the LGR5 ECD ad-infected explants became SOX9^-^/SOX2^HI^ (Figure 2B, bottom left), indicative of a loss of bud tip identity and differentiation into proximal epithelium. In addition to reduced SOX9 expression in the LGR5 ECD ad-infected explants, WNT target gene expression was reduced compared to the control, measured by *AXIN2* FISH (Figure 2C). *LGR5*, also a WNT target gene, was reduced via FISH (Figure 2D). In control ad-infected explants, *LGR5* was expressed and restricted to SOX9^+^ bud tips (Figure 2D, left panel). Although *LGR5* was still detected in the few remaining bud tip progenitors in the LGR5 ECD ad-infected explants, its endogenous expression appeared reduced relative to bud tip progenitors in control ad-infected explants, consistent with loss of bud tip progenitor identity (Figure 2D, right panel). Note that many ectopic *LGR5*^+^ cells could be detected, revealing abundant expression in strongly-infected cells, which made it difficult to quantify changes in endogenous *LGR5* expression (Figure 2D, top right). *RSPO2* expression was not affected by control or LGR5 ECD ad infection as expression was maintained at similar levels in SM22^-^ mesenchyme in both conditions (Figure 2E).

### The bud tip progenitor transcriptional profile is dependent on RSPO2-mediated signaling in bud tips

Human fetal lung explants infected with control or LGR5 ECD for 4 days were dissociated, and single cells were sequenced via scRNA-seq. Louvain clustering and UMAP visualization revealed clusters of epithelial, mesenchymal, smooth muscle, endothelial, neuroendocrine, and proliferating cells, which were identified by examining expression of canonical marker genes for these cell types (Figure S3A and S3B). There is also a cell cluster composed of only LGR5 ECD ad-infected cells, which appears to be clustered based on high expression of *LGR5* (Figure S3B and S3C, cluster 4). As a control, we sequenced non-infected explants from the same experiment (Figure S3D-F). The UMAP embedding including all three samples contained the same general cell populations, but an additional immune cell population was observed (Figure S3D and S3E, cluster 8). We noted that most clusters (with the exception of the LGR5 ECD ad-infected cluster) possessed cells from each sample that were evenly-distributed across clusters; however, part of the epithelial cell cluster showed separation between non-infected and control ad-infected cells compared to LGR5 ECD ad-infected cells (Figure S3E and S3F, cluster 2). Importantly, many mesenchymal cell populations we observed *in vivo*, including *PDGFRA*^HI^, *RSPO2*^+^, and *SM22*^+^ populations, were retained in the explants (Figure S3G).

To gain better resolution of changes occurring in the epithelium, the epithelial cell cluster (cluster 1) from the UMAP embedding that includes cells from control and LGR5 ECD ad-infected explants (Figure S3B) was extracted and re-clustered (Figure 3A). Since there appeared to be no major transcriptional differences between the control ad-infected explants and non-viral infected explants, non-infected explants were omitted from this analysis. We identified cluster 0 as bud tip progenitors based on expression of canonical bud tip progenitor marker genes such as *SOX9, TESC*, and *ETV5*. (Figure 3C). Based on visual inspection of this cluster and expression of bud tip markers, we noted that cells from each sample and gene expression were not evenly distributed across the cluster (Figure 3B arrow and 3C). To gain insights into possible differences between the control and LGR5 ECD ad-infected bud tip cells, cluster 0 was again extracted and re-clustered (Figure 3D). When re-clustered, there were 3 predicted clusters, with sub-cluster 0 primarily consisting of cells from the control ad-infected explants and sub-cluster 1 primarily consisting of cells from the LGR5 ad-infected explants (sub-cluster 2 contained both conditions) (Figure 3D-E). Bud tip progenitor genes (*SOX9, TESC, ETV5, CA2*) were more highly expressed in control ad-infected cells (Figure 3F). Moreover, individual cells from each sample were evaluated against a panel of the top 22 most differentially expressed genes from *in vivo* bud tip progenitors (see Table S1) (Holloway, Wu, *et al*., 2020; Miller *et al*., 2020), thus assigning a “bud tip progenitor cell score” to every cell (see methods). Consistent with the reduced expression of individual bud tip progenitor genes in LGR5 ECD ad-infected explants, cells from LGR5 ECD ad-infected explants were scored lower for having a bud tip progenitor identity compared to cells from control ad-infected explants (Figure 3G). Together, this data suggests that loss of endogenous RSPO2 activity during human lung development causes a reduction of bud tip progenitor gene expression.

**Figure 3.**
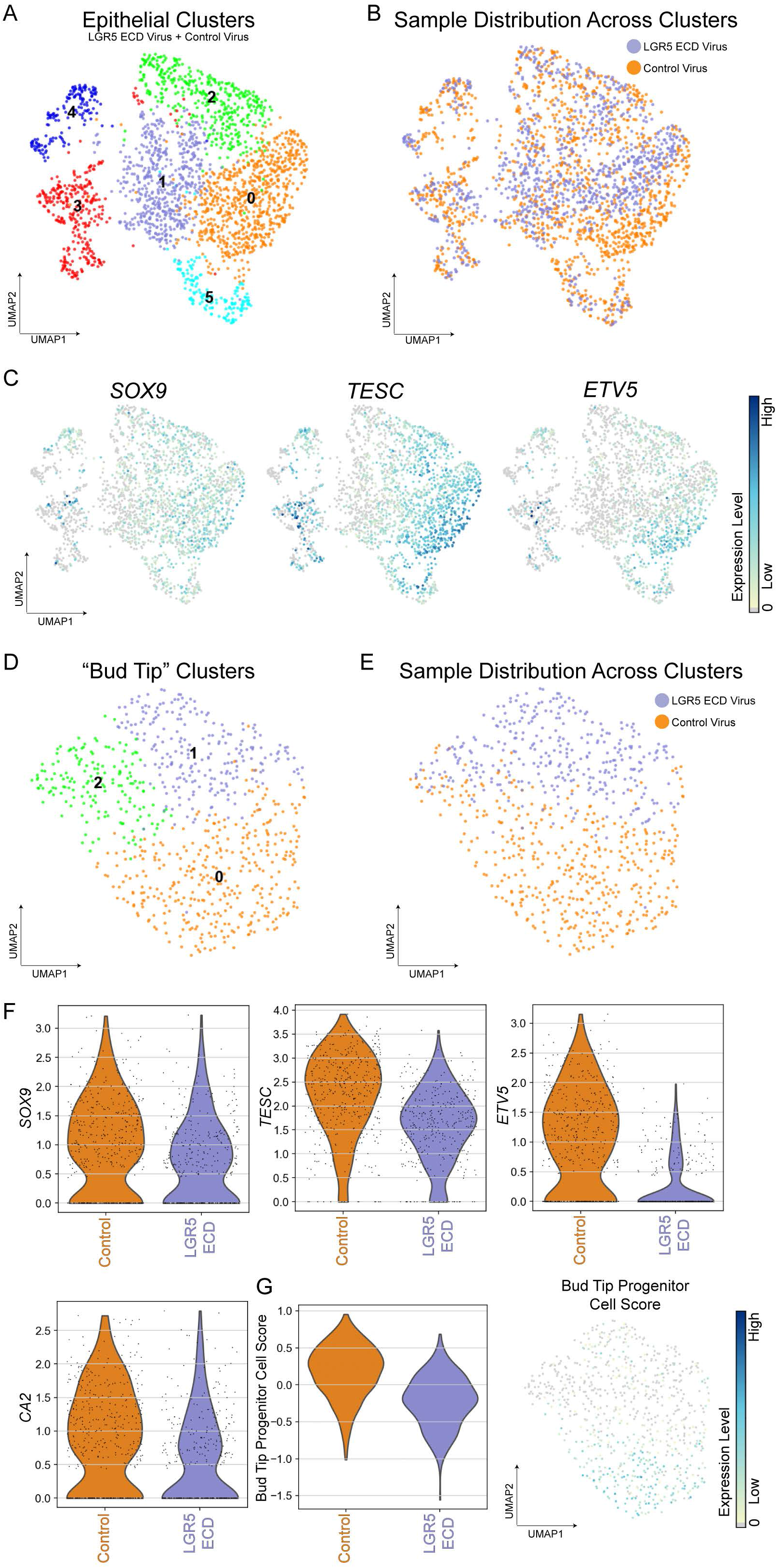
The bud tip progenitor transcriptional profile is dependent on RSPO2-mediated WNT signaling in bud tips. (A) Cluster plot of the epithelial cells (*EPCAM*^+^/*KRT18*^+^/*KRT8*^*+*^/*CLDN6*^+^) computationally extracted from LGR5 ECD adenovirus (ad)-infected explants and control ad-infected explants sequenced using single cell RNA sequencing (re-cluster of cluster 1 from Figure S3B). Each dot represents a single cell and cells were computationally clustered based on transcriptional similarities. The plot is colored and numbered by cluster. (B) UMAP plot corresponding to Figure 3A. Each dot represents a single cell and dots/cells are colored by the sample from which they came from. Cluster 0 of the cluster plot is largely separated by sample while the remaining clusters contain evenly-dispersed cells from the control ad-infected cells and LGR5 ECD ad-infected explants. (C) UMAP feature plots corresponding to the cluster plot in Figure 3A and displaying expression levels of the known bud tip progenitor markers *SOX9, TESC*, and *ETV5*. The color of each dot indicates log-normalized and z-transformed expression level of the given gene in the represented cell. The portion of cluster 0 from the UMAP plot in Figure 3A/B dominated by control ad-infected cells shows higher expression of bud tip progenitor markers. (D) Cluster plot of bud tip-like cells (re-cluster of cluster 0 from Figure 3A). Each dot represents a single cell and cells were computationally clustered based on transcriptional similarities. The plot is colored and numbered by cluster. (E) UMAP plot corresponding to Figure 3D. Each dot represents a single cell and dots/cells are colored by the sample from which they came from. The LGR5 ECD ad-infected cells and control ad-infected cells are separated where cluster 0 is composed of cells from control ad-infected explants and cluster 1 is composed of cells from LGR5 ECD ad-infected explants. Cells from both conditions compose cluster 2; however, they still remain largely separate. (F) Violin plots corresponding to the cluster plot in Figure 3D and displaying expression known of bud tip progenitor markers in control ad- and LGR5 ECD ad-infected cells. Cells from control ad-infected explants have higher expression of bud tip progenitor markers compared to cells from LGR5 ECD ad-infected explants. (G) Violin plot and UMAP feature plot corresponding to the cluster plot in Figure 3D and displaying bud tip progenitor cell score, calculated as the average expression of the top 22 enriched genes in *in vivo* bud tip progenitor cells (see methods). The color of each dot in the feature plot indicates log-normalized and z-transformed expression level of the set of bud tip genes in the represented cell. Cluster 0 in the cluster plot in Figure 3D, containing almost all control ad-infected cells and almost no LGR5 ECD ad-infected cells, shows cells with higher bud tip progenitor cell scores compared to cells in the cluster containing the LGR5 ECD ad-infected cells (cluster 1), and the violin plot shows the quantification of this.

### RSPO2-potentiated signaling in bud tips prevents differentiation into proximal cell types

Based on the reduced bud tip progenitor transcriptional profile in LGR5 ECD ad-infected explants compared to control ad-infected explants (see Figure 3), and because epithelial SOX2 expression was higher in the LGR5 ECD ad-infected explants compared to the control via IF (see Figure 2) and scRNA-seq (Figure 5A), we hypothesized that the bud tips from these explants might be differentiating into proximal lung cell types. Using scRNA-seq data from control and LGR5 ECD ad-infected explants and complementary IF and FISH stains on tissue sections from these explants, we evaluated expression of proximal and distal differentiated cell type markers. When possible, we calculated cell type scores for specific cell types by evaluating the average expression of the 50 most differentially expressed genes from *in vivo* cells of the listed cell type (Table S1) (Holloway, Wu, *et al*., 2020; Miller *et al*., 2020) (see methods). We limited the scRNA-seq analysis to cells within the bud tip cluster (see Figure 3) in order to specifically determine how the bud tips in the LGR5 ECD ad-infected cells are changing relative to controls.

The biggest difference in cell type marker expression from the control vs. LGR5 ECD ad-infected explants was expression of secretory cell genes. By FISH, the secretory cell marker *SCGB3A2* was only found in proximal airway structures in control ad-infected explants but was found in many cystic bud tip-like structures in LGR5 ECD ad-infected explants (Figure 4A). LGR5 ECD ad-infected cells had higher cell type scores for secretory progenitor and club cells but not for goblet cells compared to the control (Figure 4A). Additionally, the basal cell marker TP63 was primarily found in proximal airway structures in control ad-infected explants while expression was observed frequently in cystic, bud tip-like structures in LGR5 ECD ad-infected explants (Figure 4B). This correlated with an increase in TP63^*+*^ cell numbers in the LGR5 ECD ad-infected explants compared to the controls (Figure 4B) as well as with a higher basal cell score in cells from LGR5 ECD ad-infected explants compared to cells from control ad-infected explants (Figure 4B). Although the cell type scores for neuroendocrine and multiciliated cells were moderately increased in cells from LGR5 ECD ad-infected explants, CHGA^*+*^cells and FOXJ1^+^ cells were only sparsely detected by IF in both conditions (Figure S4A). Overall, LGR5 ECD ad-infected explants had higher expression of proximal lung lineage markers compared to control ad-infected explants, particularly with respect to secretory progenitor cell, club cell, and basal cell markers.

**Figure 4.**
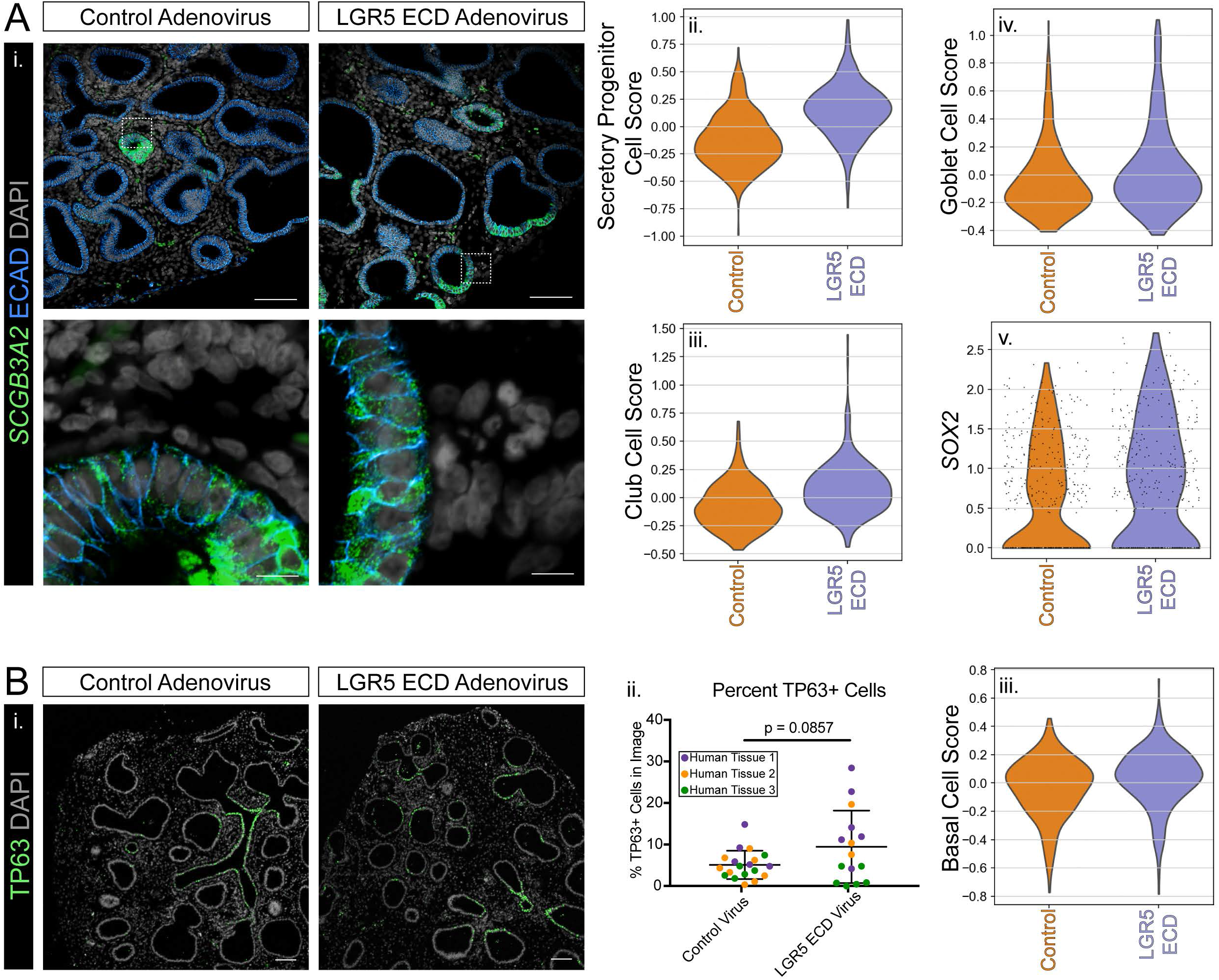
Inhibition of RSPO2-mediated WNT signaling in lung explants results in bud tip differentiation into proximal secretory and basal cell types. A) Expression of proximal secretory cell type markers in LGR5 ECD adenovirus (ad)-infected explants and control ad-infected explants. **(i)** Fluorescent *in situ* hybridization staining of the secretory cell marker *SCGB3A2* on sections from control ad-infected explants and LGR5 ECD ad-infected explants. DAPI is shown in gray. Scale bars represent 100µm or 10µm for insets. *SCGB3A2* expression is restricted to airway structures in control ad-infected explants while it is appearing in cystic bud tip structures in LGR5 ECD ad-infected explants. **(ii - iv**.) Violin plots displaying the cell score for the listed proximal secretory cell type, calculated from single cell RNA sequencing (scRNA-seq) data as the average expression of the s enriched genes in *in vivo* fetal secretory cell types (see methods). Cell score for secretory progenitor cells and club cells is higher in cells from the LGR5 ECD ad-infected explants compared to cells from the control ad-infected explants. Cell score for goblet cells is similar between the two conditions, with a small increase in the LGR5 ECD ad-infected group. **(v**.) Log-normalized and z-transformed expression level of the proximal marker *SOX2* in scRNA-seq data from LGR5 ECD ad-infected explants and control ad-infected explants. *SOX2* expression is increased in LGR5 ECD ad-infected explants compared to control ad-infected explants. B) Expression of proximal basal cell type markers in LGR5 ECD ad-infected explants and control ad-infected explants. **(i**.) Immunofluorescence staining for the basal cell marker TP63 on sections from control ad-infected explants and LGR5 ECD ad-infected explants. DAPI is shown in gray. Scale bars represent 100µm. The LGR5 ECD ad-infected explants have TP63^*+*^ cells in cystic, bud tip-like regions while TP63^*+*^ cells are primarily found in proximal airway structures in the control ad-infected explants. **(ii**.) Quantification of basal cell marker TP63 on sections from control ad-infected explants and LGR5 ECD ad-infected explants. TP63^*+*^ cells in the control ad-infected explants comprised approximately 5.6% of cells while they comprised 11.4% in LGR5 ECD ad-infected explants (p = 0.0857, Welch’s t test). This quantification was performed in three unique biological samples with one to three technical replicates and a minimum of three image fields for each sample. **(iii**.) Violin plot displaying the cell score for basal cells in LGR5 ECD and control ad-infected explants, calculated from single cell RNA sequencing data as the average expression of the top 50 enriched genes in *in vivo* fetal basal cells (see methods). Cell score for basal cells is higher in cells from the LGR5 ECD ad-infected explants compared to cells from the control ad-infected explants.

To determine if bud tip progenitors in LGR5 ECD ad-infected explants were undergoing general differentiation or differentiating specifically to proximal cell types, we examined LGR5 ECD ad-infected bud tips for the presence of distal cell types. By FISH, expression of *SFTPB* was high in all epithelium from both conditions (Figure S4B), but by scRNA-seq, was increased in LGR5 ECD ad-infected explants (Figure S4B). It has recently been shown that *SFTPB* is expressed in some proximal secretory cell types in addition to alveolar cells (Miller *et al*. 2020), which could account for higher expression in the LGR5 ECD ad-infected explants. The distal alveolar type I and type II markers ABCA3 and RAGE, respectively, were moderately increased in control explants compared to LGR5 ECD ad-infected explants at the protein level (Figure S4B), but at the transcript level, were similar or slightly increased in LGR5 ECD ad-infected explants (Figure S4B). Additional distal alveolar cell type markers were moderately increased in control ad-infected explants (*SFTPC, AQP5*) (Figure S4B) or were similarly expressed between the two conditions (*PDPN, HOPX*) (Figure S4B). It has recently been shown that the cells expressing bud tip and alveolar markers, termed bud tip adjacent cells, reside in the distal lung from approximately 12 – 20 weeks post-conception (Miller *et al*., 2020). The cell type score for bud tip adjacent cells was higher in control ad-infected explants compared to LGR5 ECD ad-infected explants (Figure S4B). Overall, there does not appear to be a strong differentiation bias into distal cell types between the two conditions, rather, we see an increase in differentiation towards proximal cell types upon LGR5 ECD ad-infection.

### Isolated RSPO2^+^ mesenchymal cells support bud tip multipotency in organoid co-cultures

To determine how the RSPO2 and SM22 mesenchymal cell populations regulate bud tip progenitor cell behavior in culture, we performed 3D co-cultures with established human fetal-derived bud tip progenitor organoids (Miller *et al*., 2018) and isolated RSPO2^+^ or SM22^+^ mesenchyme in Matrigel. We first used published approaches to identify putative cell surface markers that are co-expressed in RSPO2 cells (SurfaceGenie - (Waas *et al*., 2020)) such that we could use fluorescence active cell sorting (FACS) to separate *RSPO2*^+^ and SM22^+^ mesenchymal cells. SurfaceGenie mines scRNA-seq enrichment profiles to predict and rank cell surface candidates. The approach identified LIFR as a putative cell surface marker for specific isolation of *RSPO2*^+^ mesenchymal cells. We confirmed that *LIFR* is strongly co-expressed with *RSPO2* in mesenchymal cells prior to isolating the *RSPO2*^+^ and SM22^+^ cell populations (Figure S5A). To specifically enrich for LIFR^HI^ (*RSPO2*^+^) and LIFR^-^ (SM22^+^) mesenchyme, we used a combinatorial staining approach that allowed for the isolation of non-epithelial (EPCAM^-^), non-endothelial (CD31^-^), LIFR^HI^ or LIFR^-^ cells using FACS. (Figure S5B). RT-qPCR analysis on the isolated populations confirmed that genes enriched in *RSPO2*^*+*^ mesenchyme (*RSPO2, FGFR4*) were enriched in the LIFR^HI^ population and genes enriched in airway smooth muscle cells (*SM22, FOXF1*) were enriched in the LIFR^-^ population (Figure S5C). We also confirmed that epithelial cells and endothelial cells were also successfully depleted from the collected LIFR^HI^ and LIFR^-^ populations (Figure S5C).

Bud tips were cultured with 150,000 LIFR^HI^ or LIFR^-^ mesenchymal cells directly after FACS-isolation in a media including FGF7 and ATRA (Miller *et al*., 2018), but excluding any WNT ligands or small molecule activators. A positive control without mesenchyme but with (CHIR99021; CHIR) as well as a negative control without mesenchyme or CHIR were included in each experiment. CHIR was excluded from the co-culture media because we hypothesized that the LIFR^HI^ (RSPO2^+^) mesenchyme would provide WNT signaling cues for the bud tips. After 10 – 11 days of bud tip/mesenchyme co-culture, there were clear morphological differences in the bud tips between the LIFR^HI^ and LIFR^-^ co-cultures (Figure 5A). In both co-culture conditions, the mesenchyme caused condensation of the Matrigel and bud tips into a more tightly compacted structure compared to bud tips cultured in Matrigel without mesenchyme, but this was much more pronounced in the LIFR^-^ co-cultures (Figure 5A). Positive control (+CHIR) bud tips retained a normal, cystic phenotype while negative control (-CHIR) bud tips became dense, as expected (Figure 5A) (Miller *et al*., 2018).

**Figure 5.**
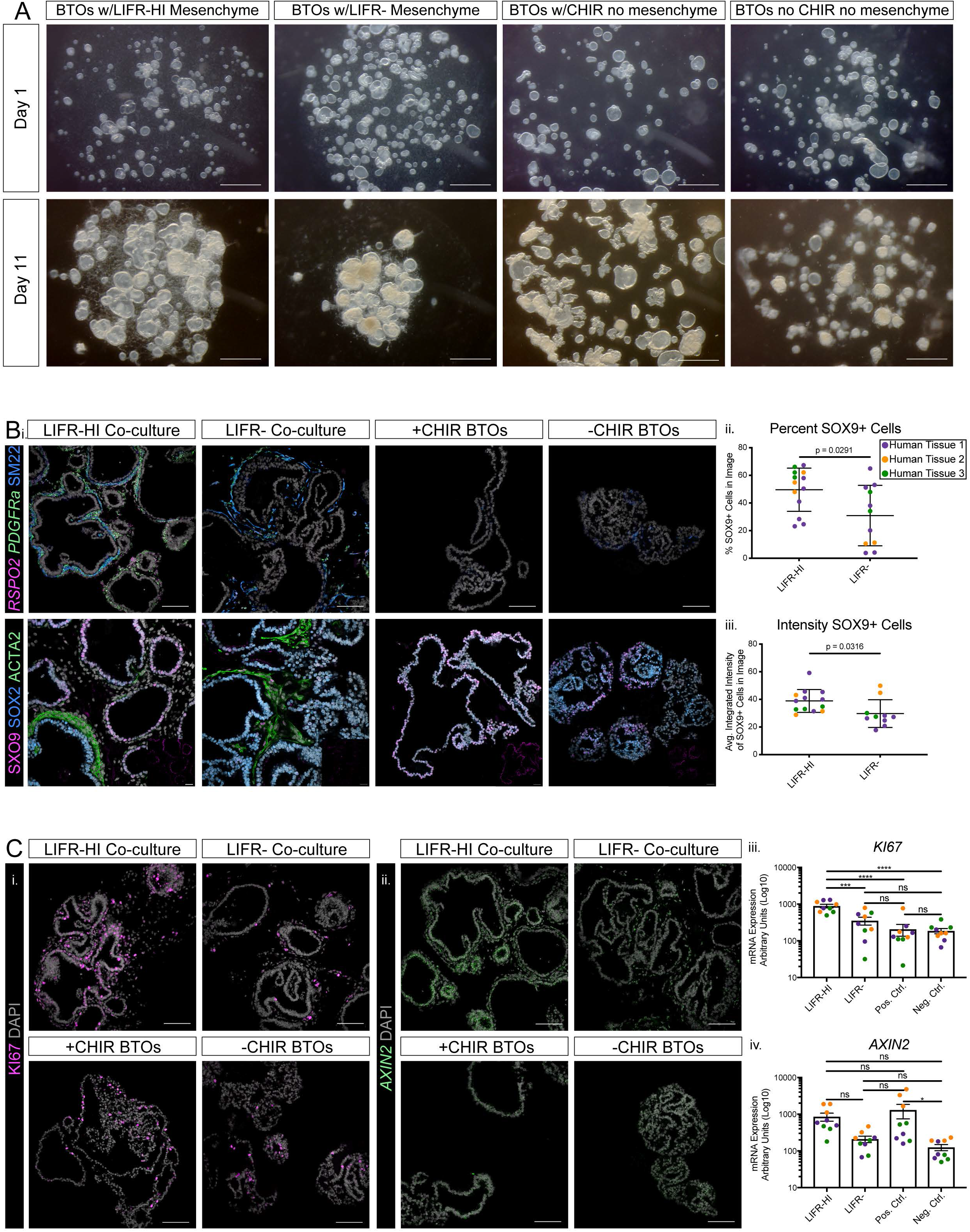
RSPO2^+^ mesenchymal cells support a proximal and distal phenotype in bud tip organoid co-cultures. **(A)** Brightfield images of human fetal lung-derived bud tip organoids co-cultured with LIFR^HI^ mesenchyme, LIFR^-^ mesenchyme, with previously-established bud tip media (Miller *et al*., 2018), or with bud tip media where CHIR99021 was removed (same media co-cultures were grown in) the day after the start of the culture (day 1) and the day of collection (day 11). Scale bars represent 1mm. In both co-culture conditions, the mesenchyme pulled the Matrigel and bud tips in more tightly compared to bud tips cultured in Matrigel without mesenchyme, but this was much more pronounced in the LIFR^-^ co-cultures. Positive control (+CHIR99021) bud tips retained a normal, cystic phenotype while negative control (-CHIR99021) bud tips became dense. **(B)** Proximal/distal epithelial patterning in relation to mesenchymal cell type localization. **(i**.) Multiplexed Fluorescence *in situ* (FISH) hybridization of *RSPO2* and *PDGFRa* and co-immunofluorescence (IF) for smooth muscle marker SM22 (top) and IF for the distal marker SOX9, proximal marker SOX2, and smooth muscle marker ACTA2 (bottom) on sections from LIFR^HI^ co-cultures, LIFR^-^ co-cultures, positive control (+CHIR99021) bud tips, and negative control (-CHIR99021) bud tips after 11 days of culture. DAPI is shown in gray. Scale bars on top panel represent 100µm. Images on the bottom panel were taken at the same magnification. Insets on the bottom panel are showing SOX9, where scale bars represent 100µm. *RSPO2*^*+*^ cells and SM22^+^ cells were found in the LIFR^HI^ co-culture while only SM22^+^ cells were found in the LIFR^-^ co-culture. *PDGFRa* expression was found throughout the mesenchyme in both co-cultures. Epithelium surrounded by ACTA2^+^ cells in the LIFR^HI^ co-culture was negative for, or had low expression of, the distal marker SOX9 and had high expression of the proximal marker SOX2 while epithelial cells surrounded by *RSPO2*^*+*^ cells retained the distal marker SOX9. Epithelium in the LIFR^-^ co-cultures was SOX^HI^ and SOX9^-^ or SOX9^LOW^. SOX9 protein expression remained high in bud tips cultured with CHIR while it was lower and expressed in fewer cells in cultures where CHIR was removed. **(ii – iii**.) Quantification of SOX9^+^ cell numbers and integrated staining intensity of SOX9 from IF stains. LIFR^HI^ co-cultures retained higher numbers of SOX9^*+*^ cells (p = 0.0291, Welch’s t test) at higher levels (p = 0.0316, Welch’s t test) compared to LIFR^-^ co-cultures. A) KI67 and *AXIN2* staining and quantification. **(i. – ii**.) IF for KI67 (left) and FISH for *AXIN2* (right) on sections from LIFR^HI^ co-cultures, LIFR^-^ co-cultures, positive control (+CHIR99021) bud tips, and negative control (-CHIR99021) bud tips after 11 days of culture. DAPI is shown in gray. Scale bars represent 100µm. KI67^+^ cells and *AXIN2* expression were detected in each condition but both appear highest in the LIFR^HI^ co-culture. **(iii. – iv**.) RT-qPCR for *KI67* and *AXIN2* on LIFR^HI^ co-cultures, LIFR^-^ co-cultures, positive control (+CHIR99021) bud tips, and negative control (-CHIR99021) bud tips after 11 days of culture from three independent experiments. *KI67* expression is significantly higher in the LIFR^HI^ co-culture compared to all other conditions. *AXIN2* expression is highest in the positive control, followed by the LIFR^HI^ co-culture, then the LIFR^-^ co-culture, and finally the negative control. Each color represents an independent experiment using bud tips and mesenchyme from unique specimens. Each data point of the same color represents a technical replicate from the same set of tissue specimens. Error bars represent standard error of the mean. Statistical tests were performed by ordinary one-way ANOVA followed by Dunnett’s multiple comparison test. ‘*’ represents a p-value less than 0.05, ‘**’ represents a p-value less than 0.01, ‘***’ represents a p-value less than 0.001, ‘****’ represents a p-value less than 0.0001, and ‘ns’ represents a p-value above 0.05.

By FISH, *RSPO2*^*+*^ cells were found in the LIFR^HI^ co-culture after 11 days of culture (Figure 5B). Surprisingly, SM22^+^ cells were also detected in the LIFR^HI^ co-culture at this time point, but at lower numbers and expression levels compared to the LIFR^-^ co-culture (Figure 5B). Within LIFR^HI^ co-cultures, *RSPO2* expression was absent from SM22^+^ cells, except for some SM22^LOW^ cells (Figure 5B), while *RSPO2* expression was absent from the LIFR^-^ co-culture. *PDGFRA* expression was found throughout the mesenchyme in both co-cultures (Figure 5B). By IF, epithelial cells surrounded by ACTA2^+^ (another marker for smooth muscle) cells in the LIFR^HI^ co-culture were negative for, or had low expression of, the distal marker SOX9 and had high expression of the proximal marker SOX2, while epithelial cells surrounded by *RSPO2*^*+*^ cells retained the distal marker SOX9 (Figure 5B). This data suggests that both proximal and distal epithelium can differentiate when bud tips are co-cultured with LIFR^HI^ cells. Epithelium in the LIFR^-^ co-cultures was SOX2^HI^ and SOX9^- or LOW^ (Figure 5B), indicative of epithelium undergoing proximal differentiation. In controls, SOX9 protein expression remained high in bud tips cultured with CHIR while it was much lower and expressed in fewer cells in cultures where CHIR was removed (Figure 5B).

Bud tip genes *SOX9, ETV5, NPC2*, and *LGR5* were increased in the LIFR^HI^ co-cultures compared to the LIFR^-^ co-cultures by RT-qPCR, although not by a statistically significant level (Figure S6B). Given that the LIFR^HI^ co-cultures had both SOX2^+^/SOX9^-^ proximal and SOX2^+^/SOX9^+^ distal phenotypes in the epithelium, we decided to investigate specific proximal and distal differentiation markers. The proximal basal cell marker TP63 was detected in many cells by IF in all conditions except for the positive control (Figure S6A). Additionally, proximal secretory, basal, goblet, and multicilited cell genes were highly expressed in the LIFR^HI^ co-culture, LIFR^-^ co-culture, and negative control (-CHIR) compared to the (+CHIR) positive control by RT-qPCR (Figure S6B). Interestingly, the distal differentiation/alveolar genes *SFTPC, ABCA3*, and *RAGE* were significantly up-regulated in the LIFR^HI^ co-culture compared to the LIFR^-^ co-culture and the negative control (Figure S6B). At the protein level, SFTPC was detected at high levels in the LIFR^HI^ co-culture and was low to absent in all other conditions (Figure S6A). Some SFTPC^+^ cells in the LIFR^HI^ co-culture co-expressed the other alveolar type II proteins, including SFTPB and HTII-280 (Figure S6A), and the alveolar type I marker RAGE was also detected in the LIFR^HI^ co-culture (Figure S6A). Abundant SFTPB^+^ cells were detected in the LIFR^-^ co-culture and in the negative control, but they co-expressed *SCGB3A2* (Figure S6A), indicative of a proximal differentiation (Miller *et al*., 2020). Although *SCGB3A2* mRNA was detected in the LIFR^HI^ co-culture and the positive control, protein expression was low in the LIFR^HI^ co-culture and almost absent from the positive control (Figure S6A). Together, this data suggests that LIFR^HI^/RSPO2^+^ mesenchymal cells support the multipotency of bud tip progenitors to give rise to proximal and distal cell types, while smooth muscle (LIFR^-^/SM22^+^) cells only support proximal differentiation of bud tips.

In addition to differentiation, we wanted to determine if there were changes in cell proliferation between the two co-culture conditions. We found that KI67 expression was significantly up-regulated in the LIFR^HI^ co-culture compared to the LIFR^-^ co-culture as well as compared to the positive and negative control (Figure 5C). This correlated with higher *AXIN2* expression in the LIFR^HI^ co-culture compared to the LIFR^-^ co-culture and negative control, though this was a trend and not a statistically significant change (Figure 5C). This data suggests that RSPO2^+^ mesenchymal cells support a high WNT signaling niche conducive for self-renewal (proliferation) and differentiation of bud tip progenitor cells into both proximal and distal airway.

## DISCUSSION

The lung development field has well-established literature interrogating the diversity of cell types and functions in the epithelium; however, less is known with respect to the developing lung mesenchyme, and particularly in the context of the human lung. Most of what is known about the role of the lung mesenchyme during development has come from animal models and has primarily focused on understanding airway smooth muscle cells or mesenchymal heterogeneity and function during alveolar and later stages of development (Torday, Torres and Rehan, 2003; Chen *et al*., 2012; McQualter *et al*., 2013; El Agha and Bellusci, 2014; Li *et al*., 2015, 2018, 2020; Green *et al*., 2016; Zepp *et al*., 2017; Endale *et al*., 2017; Kishimoto *et al*., 2018; Wu *et al*., 2018; Noe *et al*., 2019; Goodwin *et al*., 2019, 2020; Guo *et al*., 2019; Han *et al*., 2019; Bridges *et al*., 2020; Riccetti *et al*., 2020; Yin and Ornitz, 2020; Gouveia *et al*., 2020; Negretti *et al*., 2021). There are also far fewer similar studies in humans (Rehan *et al*., 2006; Danopoulos, Shiosaki and Al Alam, 2019; Du *et al*., 2019; Goodwin *et al*., 2019; Leeman *et al*., 2019; Shiraishi, Nakajima, *et al*., 2019; Shiraishi, Shichino, *et al*., 2019; Danopoulos *et al*., 2020; Guney *et al*., 2020). Nevertheless, recent advances in single cell analytical tools and *in vitro* human-specific model systems have allowed us to begin addressing unknowns in human lung development (Treutlein *et al*., 2014; Brazovskaja, Treutlein and Camp, 2019; Du *et al*., 2019; Kishimoto *et al*., 2019; Travaglini *et al*., 2019; Danopoulos *et al*., 2020; Miller *et al*., 2020; Yu *et al*., 2020). Here, we aimed to understand how mesenchymal cells are involved in creating an epithelial bud tip progenitor cell niche.

Single cell sequencing analysis predicted airway smooth muscle cells and three non-smooth muscle populations that have similar but not identical gene expression profiles in the developing human lung. We observed that the non-smooth muscle mesenchymal cells identified by scRNA-seq express the WNT-agonist *RSPO2* throughout the developmental time frame analyzed, and spatial localization showed that *RSPO2* is expressed adjacent to the bud tip domain. It is known that high WNT signaling conditions are necessary for the maintenance of the bud tip progenitor cell state *in vitro* (Nikolić *et al*., 2017; Miller *et al*., 2018; Rabata *et al*., 2020), which made this cell population a strong bud tip-associated mesenchymal cell candidate. By using human tissue specimens and human *in vitro* model systems to explore this cell population further, we provide evidence that the *RSPO2*^*+*^ mesenchymal cells signal to bud tip progenitors, likely through LGR5, in order to maintain a high WNT signaling zone in the bud tips. We show that this RSPO2-mediated WNT signaling niche provides support for the bud tips to maintain their progenitor state and give rise to both distal alveolar cell types as well as proximal airway cell types. The process of RSPO ligands signaling through LGR receptors to maintain high WNT signaling in progenitor and stem cells has been described in other organs and tissues, such as in the intestinal crypt, skin, hair follicle, and mammary tissue (Barker *et al*., 2007; Jaks *et al*., 2008; Trejo *et al*., 2017; Yan *et al*., 2017; Dame *et al*., 2018; Baulies, Angelis and Li, 2020; Holloway, Czerwinski, *et al*., 2020). However, an alternative mechanism where RSPO2 acts independently of LGRs, and instead through RNF43/ZNRF3, has been described in the context of limb development (Szenker-Ravi *et al*., 2018). It is possible that blocking endogenous RSPO2 through the LGR5 ECD adenovirus is preventing RSPO2 from binding these alternative receptors. Appropriate genetic loss-of-function experiments to formally test this hypothesis is difficult-to-impossible in human tissue. Through human fetal lung explants infected with an LGR5 ectodomain adenovirus that disrupts the RSPO2-mediated WNT signaling axis, we show that bud tips lose their identity and differentiate into proximal cell types. Our data suggests a potential differentiation bias towards secretory progenitor, club, and basal cells; however, previous data using cultured bud tip progenitors shows that removal of the WNT component of the media causes an up-regulation of secretory cell markers, suggesting secretory cells may just be a default differentiation state (Miller *et al*., 2018). Moreover, in mice, multiciliated, goblet, and neuroendocrine cells appear later in development compared to secretory and basal cell types (Rock *et al*., 2009; Treutlein *et al*., 2014; Ardini-Poleske *et al*., 2017; Montoro *et al*., 2018; Miller *et al*., 2020).

Through mesenchyme and bud tip organoid co-cultures, we show that *RSPO2*^*+*^ mesenchymal cells allow bud tips to give rise to distal alveolar cell types as well as proximal airway cell types. Although we successfully separated *RSPO2*^*+*^ mesenchymal cells from SM22^*+*^ mesenchymal cells via FACS prior to co-culture experiments, we still detected many SM22^*+*^ mesenchymal cells in the FACS-isolated *RSPO2*^*+*^ mesenchyme and bud tip co-cultures after the 10 – 11-day culture period. The areas where proximal differentiation occurred in the *RSPO2*^*+*^ mesenchyme and bud tip co-cultures correlated to the areas where SM22^*+*^ mesenchymal cells were. It is possible that enough SM22^*+*^ mesenchymal cells were captured in the LIFR^HI^ (RSPO2^+^) FACS-isolated population due to sorting error and were able to expand over the culture period. However, it is also possible that RSPO2^*+*^ mesenchymal cells give rise to SM22^*+*^ airway smooth muscle cells, which support proximal differentiation. This idea is especially supported by the fact that SM22^LOW^ cells in the LIFR^HI^ co-cultures expressed low levels of *RSPO2*. There is much to be learned about the intricate dynamics of the mesenchyme during human lung development.

Of particular interest for the current study, *RSPO2* mutations in humans are lethal at birth, causing nearly complete lung aplasia with lung development ceasing just after the primary lung buds emerge from the trachea (Szenker-Ravi *et al*., 2018), supporting a critical role for *RSPO2*^*+*^ mesenchymal cells during human lung development. Although non-functional *Rspo2* in mice is also lethal at birth, with mutant lungs exhibiting reduced branching, laryngeal-tracheal defects, and reduced Wnt signaling in the bud tips, mutant lungs undergo branching and do not show nearly as severe lung aplasia seen in humans (Bell *et al*., 2008). There are multiple possible explanations for why humans and mice have different severities of developmental defects caused by *RSPO2* mutations. First, it is possible that *Rspo1, Rspo3*, or *Rspo4* can compensate for the loss of *Rspo2* in mice, which may not exist in humans. Our scRNA-seq and FISH data for *RSPO1, RSPO3*, and *RSPO4* in the human distal lung indicate that they are expressed in the same broad cell population as *RSPO2*, but at much lower levels. It would be valuable to determine the expression patterns of the other *Rspo* transcripts and proteins in the murine lung and determine if other RSPOs can replace the role of RSPO2 in the murine and human lungs. The advent of *in vitro* model systems of the developing human lung provides an excellent opportunity to explore these questions further (Conway *et al*., 2020).

Another possibility for the mouse/human phenotype difference is if *RSPO2* is necessary for initiating branching morphogenesis during the earliest stages of human, but not mouse, lung development. In mice, deletion of *Wnt2* and *Wnt2b* together inhibit lung progenitors from ever being specified, and deletion of *Wnt2* and *Wnt7b* together result in defective branching and non-localized SOX9 expression (Goss *et al*., 2009; Miller *et al*., 2012). The possible necessity of *RSPO2* to promote WNT signaling that may be necessary for maintaining lung progenitors, and preventing precocious proximal differentiation, in the primary lung buds and/or to initiate branching morphogenesis in humans could explain why lungs in humans lacking functional *RSPO2* fail to develop past the primary lung bud stage. Through infection of human fetal lung explants with an LGR5 ECD adenovirus that reduces RSPO2-mediated WNT signaling, we show that premature differentiation of bud tip progenitors into proximal lineages occurs. Although we show this phenomenon after branching morphogenesis has already begun, because of *RSPO2*’s persistence throughout all the time points sequenced in this study, this could be the case beginning at the primary lung buds. Therefore, if premature differentiation of lung progenitors occurs at the primary lung bud stage, the lung could fail to develop further.

The findings in this study have also prompted additional questions. It is known that WNT signaling is an important regulator of human bud tip progenitor maintenance (Miller *et al*., 2018); however, for the first time, we can appreciate the much larger signaling network involved in maintaining WNT signaling in the bud tip niche. How other cell types and signaling pathways may be integrated to control bud tip progenitor behavior is a fascinating avenue of future exploration. Additionally, although *RSPO2*^*+*^ mesenchymal cells are localized adjacent to bud tip progenitor cells, expression of *RSPO2* extends far beyond the cells that sit near bud tip progenitors. Combined with the unique expression pattern of *LGR4* throughout the mesenchyme and *LGR6* in airway smooth muscle cells, it would be interesting to understand the role that *RSPO2*^*+*^ cells have a role in regulating the behavior of other mesenchymal cells. The role of RSPOs may also extend beyond the distal lung and into the proximal lung. Overall, the current study reveals that *RSPO2*^+^ cells form the bud tip progenitor niche in the developing human lung and opens up many questions for further understanding the complex process of human lung development.

## ACKNOLEDGMENTS

Financial support: This work was supported by a Chan Zuckerberg Initiative Human Cell Atlas Seed Network grant and by the NIH-NHLBI (R01HL119215) funding to J.R.S. R.F.C.H. was supported by a NIH Tissue Engineering and Regenerative Medicine Training Grant (NIH-NIDCR T32DE007057) and by a Ruth L. Kirschstein Predoctoral Individual National Research Service Award (NIH-NHLBI F31HL152531). A.J.M. was supported by a Ruth L. Kirschstein Predoctoral Individual National Research Service Award (NIH-NHLBI F31HL142197). E.M.H. was supported by a Ruth L. Kirschstein Predoctoral Individual National Research Service Award (NIH-NHBLI F31HL146162). T.F. was supported by a NIH Tissue Engineering and Regenerative Medicine Training Grant (NIH-NIDCR T32DE007057). A.S.C. was supported by the T32 Michigan Medical Scientist Training Program (5T32GM007863-40).

We would like to thank Judy Opp and the University of Michigan Advanced Genomics core for the operation of the 10X Chromium single cell capture platform, the University of Michigan Microscopy core for providing access to confocal microscopes, the Flow Cytometry core for providing access to flow cytometers, and the Vector core for providing adenovirus purification and expansion services. We would also like to thank the University of Washington Laboratory of Developmental Biology.

## AUTHOR CONTRIBUTIONS

R.F.C.H., J.R.S., E.S.-R., and B.R. conceived the study. J.R.S. supervised the research. R.F.C.H. designed, performed, and interpreted experiments characterizing human fetal lung mesenchyme and utilizing *in vitro* human fetal lung explant cultures and co-cultures to functionally assess fetal lung mesenchyme populations. Y.-H.T., A.W., E.M.H., and A.J.M. developed and executed tissue dissociation methods for scRNA-seq, J.H.W. performed computational analysis on scRNA-seq data, and R.F.C.H. and J.R.S. interpreted scRNA-seq results. R.F.C.H, Y.-H.T., A.W., E.M.H., and A.J.M. processed fetal lung tissues for IF/FISH. R.F.C.H. and E.M.H. performed FACS experiments. R.F.C.H., A.J.M., T.F., and A.S.C. derived human fetal bud tip cultures. R.F.C.H., A.J.M., T.F., and A.S.C. designed human fetal lung explant culture systems. K.S.Y. and C.J.K. developed and provided the LGR5 ECD and control adenovirus. T.F. and A.S.C. provided insight into data interpretation. R.F.C.H. assembled figures, and and J.R.S. wrote the manuscript. J.H.W. contributed to writing the methods. All authors read, contributed feedback, and approved the manuscript.

## DECLARATION OF INTERESTS

The authors have no competing interests.

## METHODS

### Lead Contact

Please contact Jason R. Spence at if you would like to request materials used in this study.

## Materials Availability

This study did not generate any new reagents.

## Data and Code Availability

Sequencing data used in this study is deposited at EMBL-EBI ArrayExpress. Single-cell RNA sequencing of human fetal lung and human fetal lung explants: human fetal lung (ArrayExpress: E-MTAB-8221) (Miller *et al*., 2020), human fetal lung explants (ArrayExpress: in progress) (this study). Code used to process data can be found at: https://github.com/jason-spence-lab/Hein_2021.

## Experimental Models and Subject Details

### Human Lung Tissue

Research involving human lung tissue (8.5 – 19 weeks post conception) was approved by the University of Michigan Institutional Review Board. All human lung tissue used in these experiments was normal, de-identified tissue obtained from the University of Washington Laboratory of Developmental Biology. The tissue was shipped overnight in Belzer-UW Cold Storage Solution (Thermo Fisher, NC0952695) on ice, and all experiments were performed within 24 hours in Belzer solution.

### Lung Explants

Three unique human tissue samples spanning 11 – 13 weeks post-conception were used. For each unique tissue, one to three explants were included for each type of analysis.

#### Culture Establishment

For air-liquid-interface culture, Nucleopore Track-Etched Membranes (13mm, 8µm pore, polycarbonate) (Sigma, Cat#WHA110414) placed in 24-well tissue culture plates (Thermo Fisher, Cat#12-565-163) were pre-coated with 20µg/cm^2^ Collagen Type I (Sigma, Cat#C5533) in 0.01N ice-cold acetic acid for 30 minutes on ice followed by 2 hours at 37°C. The membranes were then washed with 1X PBS directly before use. To prepare the explants, the lung was placed in a petri dish in ice-cold 1X PBS and approximately 1mm^2^ pieces of tissue were cut from the most distal edge of the lung under a stereomicroscope using forceps and a scalpel. 500mL culture media containing Advanced DMEM/F-12 (Thermo Fisher, Cat#12634010), 100µg/mL penicillin-streptomycin (Thermo Fisher, Cat#15140122), 2mM L-Glutamine (Thermo Fisher, Cat#25030081), 10mM HEPES (Corning, Cat#25060CI), 1 bottle B-27 Supplement (Thermo Fisher, Cat#17504044), 1 bottle N-2 Supplement (Thermo Fisher, Cat#17502048), and 0.4µM 1-Thioglycerol (Sigma, Cat#M1753) was added to each well in the plate underneath the membrane. One explant per membrane was placed directly on the membrane in the center. Media was changed every 2 days.

#### Infection with adenovirus

Immediately following placement of the explants on the Nucleopore Track-Etched Membranes, 10^10^pfu of control or LGR5 ECD adenovirus previously described in Yan *et al*. (Yan *et al*., 2017) was pipetted directly on top of each explant under a stereo microscope using a p10 pipette. A maximum of 2µL adenovirus was added to each explant at a time to prevent the adenovirus from running off the explant. The explants were re-infected every 2-3 days. Infection of the tissue was confirmed by immunofluorescence for FLAG and the murine IgG2a Fc fragment for the LGR5 ECD adenovirus and control adenovirus respectively.

### Bud Tip Organoid and Mesenchyme Co-Cultures

#### Establishment of Bud Tip Organoid Lines

Human fetal lung bud tip organoids were derived as previously reported (Miller *et al*., 2018). In short, the lung was placed in a petri dish in ice-cold 1X PBS, and approximately 1cm^2^ pieces of tissue were cut from the most distal edge of the lung under a stereomicroscope using forceps and a scalpel. The tissue was enzymatically digested using dispase (Corning, Cat#354235) for 30 minutes then 100% FBS (Sigma, Cat#12103C) for 15 minutes. In DMEM/F-12 (Corning, Cat#10-092-CV) supplemented with 10% FBS and 100µg/mL penicillin-streptomycin (Thermo Fisher, Cat#15140122), the tissue was vigorously pipetted with a P1000 and subsequently a P200 to dissociate the epithelium from the mesenchyme, then was washed multiple times in 1X PBS to obtain as pure a population of epithelial bud tips as possible. Bud tips were plated in ∼20µL 8mg/mL Matrigel (Corning, Cat#354234) droplets in 24-well tissue culture plates (Thermo Fisher, Cat#12-565-163) and fed every 3-4 days in previously published bud tip media (Miller *et al*. 2018). Stable and epithelial-only bud tip organoids were established through a minimum of one passage before experiments began. All experiments involving bud tip cultures were derived from lungs 16.5 – 18 weeks post-conception.

#### FACS of Mesenchymal Cells

The lung (10 – 11.5 weeks post-conception) was placed in a petri dish and approximately 1 gram was cut from the most distal edge using forceps and a scalpel. The tissue was minced as much as possible using dissecting scissors, then was placed into a 15mL conical tube containing 9mL 0.1% (w/v) filter-sterilized Collagenase Type II (Thermo Fisher, Cat#17101015) in 1X PBS and 1mL filter-sterilized 2.5 units/mL dispase (Thermo Fisher, Cat#17105041) in PBS. The tube was placed at 37°C for 60 minutes with mechanical disruption using a serological pipette every 10 minutes. After 30 minutes, 75µL DNase I was added to the tube. 5mL isolation media containing 78% RPMI 1640 (Thermo Fisher, Cat#11875093), 20% FBS (Sigma, Cat#12103C), 100µg/mL penicillin-streptomycin (Thermo Fisher, Cat#15140122), and 2mM L-Glutamine (Thermo Fisher, Cat#25030081) was added. Cells were passed through 100µm and 70µm cell strainers, pre-coated with isolation media, and centrifuged at 400g for 5 minutes at 4°C. 1-2mL Red Blood Cell Lysis Buffer (Sigma, Cat#11814389001) and 0.5-1mL FACS buffer (2% BSA, 10µM Y-27632 (APExBIO, Cat#A3008), 100µg/mL penicillin-streptomycin) was added to the tube, and the tube was rocked for 15 minutes at 4°C. The cells were centrifuged at 500g for 5 minutes at 4°C, washed twice in 2mL FACS buffer, re-suspended in FACS buffer, and counted. 10^6^ cells were placed into FACS tubes (Corning, Cat#352063) for all control tubes (no antibody, DAPI only, isotype controls, individual antibodies/fluorophores) and 8 × 10^6^ cells were placed into a FACS tube for cell sorting. Primary antibodies were added at room temperature (30 minutes for LIFR and corresponding isotype, 10 minutes for CD324 and CD31 and corresponding isotypes) (see Table S2 for antibody dilutions). 3mL FACS buffer was added to each tube, then tubes were centrifuged at 300g for 5 minutes at 4°C. Cells were washed twice with 3mL FACS buffer, centrifuging at 300g for 5 minutes at 4°C between washes. Cells were resuspended in FACS buffer and 0.2µg/mL DAPI was added to appropriate tubes. FACS was performed using a Sony MA900 cell sorter and accompanying software. LIFR^HI^/CD324^-^/CD31^-^ cells and LIFR^-^ /CD324^-^/CD31^-^cells were collected in 1mL isolation media. LIFR^HI^ cells were gated highest 30% of LIFR expression.

#### Bud Tip Organoid and Mesenchyme Co-cultures

Established bud tip organoids were placed into a microcentrifuge tube and removed from Matrigel by pipetting with a P1000. Bud tips that had not been passaged within 10 days were also passed 1x through a 27-guage needle. The bud tips were then centrifuged for ∼10 seconds in a microcentrifuge and the media and Matrigel was removed under a stereomicroscope. Freshly FACS-isolated mesenchymal cells were immediately counted using a hemocytometer and enough cells were pelleted to reach approximately 150,000 mesenchymal cells per well. Matrigel (Corning, Cat#354234) was added to the tubes containing bud tip organoids on ice. For co-cultures, the bud tips in Matrigel were transferred to the tubes containing the mesenchymal cell pellets. The bud tip organoids and mesenchyme were thoroughly mixed in the Matrigel by pipetting and swirling a P200 with the tip cut off. ∼20µL droplets of Matrigel with bud tip organoids +/- mesenchyme were placed into the center of wells of a 24-well tissue culture plate (Thermo Fisher, Cat#12-565-163). The plate was inverted and placed in an incubator at 37°C for 20 minutes. For co-culture and negative control wells, 0.5mL media consisting of DMEM/F-12 (Corning, Cat#10-092-CV), 100µg/mL penicillin-streptomycin (Thermo Fisher, Cat#15140122), 2mM L-Glutamine (Thermo Fisher, Cat#25030081), 1 bottle B-27 supplement (Thermo Fisher, Cat#17504044), 1 bottle N-2 supplement (Thermo Fisher, Cat#17502048), 0.05% BSA (Sigma, Cat#A9647) and final concentrations of 50µg/mL L-ascorbic acid (Sigma, Cat#A4544), 0.4µM 1-Thioglycerol (Sigma, Cat#M1753), 50nM all trans retinoic acid (Sigma, Cat#R2625), and 10ng/mL recombinant human FGF7 (R&D Systems, Cat#251-KG) were added to each well. For positive control wells, 3µM CHIR99021 (STEMCELL Technologies, Cat#72054) was added to the above media. The cultures were fed every 3-4 days and were cultured for a total of 10-11 days. Each biological replicate shown is from a unique bud tip line co-cultured with mesenchymal cells from a unique human tissue specimen, and all three experiments were performed with four technical replicates.

## Method Details

### scRNA-seq Tissue Processing

All tubes and pipette tips were pre-washed with 1% BSA in 1X HBSS (all HBSS in protocol is with Mg^2+^ and Ca^2+^) to prevent cell adhesion to the plastic. The tissue was placed in a petri dish in ice-cold 1X HBSS, and the tissue was minced under a stereomicroscope using scissors. For uncultured lung tissue, roughly 1cm^2^ of the most distal portion of the lung was isolated, and for lung explants, 4 explants were collected per condition. The minced tissue was transferred to a 15mL conical tube with the HBSS, centrifuged at 500g for 5 minutes at 10°C, and the HBSS was removed. Mix 1 from the Neural Tissue Dissociation Kit (Miltenyi, Cat#130-092-628) was added to each tube, and the tube was placed at 37°C for 15 minutes, then Mix 2 was added and the cells were incubated for 10 minutes at 37°C. The cells were agitated by harshly pipetting with a P1000. The incubation/agitation step was repeated every 10 minutes until the cells looked to be a single cell suspension, approximately 30 minutes. The cells were then filtered through a 70µm filter, pre-coated with 1% BSA in 1X HBSS, into a 15mL conical tube. The filter was rinsed 3x with 1mL 1% BSA in 1X HBSS. The cells were centrifuged at 500g for 5 minutes at 10°C, and the supernatant was removed. 1mL Red Blood Cell Lysis Buffer (Sigma, Cat#11814389001) and 0.5mL 1% BSA in 1X HBSS was added to the tube, and the tube was rocked for 15 minutes at 4°C. The cells were centrifuged at 500g for 5 minutes at 10°C, washed twice in 2mL 1% BSA in 1X HBSS and centrifuged again. The cells were resuspended in 200µL 1% BSA in 1X HBSS, counted using a hemocytometer, centrifuged at 500g for 5 minutes at 10°C, and resuspended to reach a concentration of 1,000 cells/µL. Approximately 100,000 cells were put on ice and single cell libraries were immediately prepared on the 10x Chromium at the University of Michigan Sequencing Core with a target of 10,000 cells.

### scRNA-seq Quantification and Statistical Analysis

#### Overview

To visualize distinct cell populations within the single cell RNA sequencing dataset, we employed the general workflow outlined by the Scanpy Python package (Wolf, Angerer and Theis, 2018). This pipeline includes the following steps: filtering cells for quality control, log normalization of counts per cell, extraction of highly variable genes, regressing out specified variables, scaling, reducing dimensionality with principal component analysis (PCA) and uniform manifold approximation and projection (UMAP) (McInnes, Healy and Melville, 2018), and clustering by the Louvain algorithm (Blondel *et al*., 2008).

#### Sequencing data and processing FASTQ reads into gene expression matrices

All single-cell RNA sequencing was performed at the University of Michigan Advanced Genomics Core with an Illumina Novaseq 6000. The 10x Genomics Cell Ranger v# pipeline was used to process raw Illumina base calls (BCLs) into gene expression matrices. BCL files were demultiplexed to trim adaptor sequences and unique molecular identifiers (UMIs) from reads. Each sample was then aligned to the human reference genome (hg19) to create a filtered feature bar code matrix that contains only the detectable genes for each sample.

#### Quality Control

To ensure quality of the data, all samples were filtered to remove cells expressing too few or too many genes (Figure 1, S1, S5 - <750, >3000; Figure 3, 4, S3, S4 - <500, >10000) with high UMI counts (Figure 1, S1, S5 - >15000; Figure 3, 4, S3, S4 - >50000), or a fraction of mitochondrial genes greater than (Figure 1, S1, S5 - >0.05, Figure 3, 4, S3, S4 - >0.1)

#### Normalization and Scaling

Data matrix read counts per cell were log normalized, and highly variable genes were extracted. Using Scanpy’s simple linear regression functionality, the effects of total reads per cell and mitochondrial transcript fraction were removed. The output was then scaled by a z-transformation.

#### Variable Gene Selection

Highly variable genes were selected by splitting genes into 20 equal-width bins based on log normalized mean expression. Normalized variance-to-mean dispersion values were calculated for each bin. Genes with log normalized mean expression levels between 0.125 and 3 and normalized dispersion values above 0.5 were considered highly variable and extracted for downstream analysis.

#### Batch Correction

We have noticed batch effects when clustering data due to technical artifacts such as timing of data acquisition or differences in dissociation protocol. To mitigate these effects, we used the Python package BBKNN (batch balanced k nearest neighbors) (Polański *et al*., 2019). BBKNN was selected over other batch correction algorithms due to its compatibility with Scanpy and optimal scaling with large datasets. This tool was used in place of Scanpy’s nearest neighbor embedding functionality. BBKNN uses a modified procedure to the *k*nearest neighbors’ algorithm by first splitting the dataset into batches defined by technical artifacts. For each cell, the nearest neighbors are then computed independently per batch rather than finding the nearest neighbors for each cell in the entire dataset. This helps to form connections between similar cells in different batches without altering the PCA space. After completion of batch correction, cell clustering should no longer be driven by technical artifacts.

#### Dimension Reduction and Clustering

Principal component analysis (PCA) was conducted on the filtered expression matrix followed. Using the top principal components (Figure 1, S5 – 10; Figure 3, 4, S4 – 11; Figure S1 – 20; Figure S3 – 15), a neighborhood graph was calculated for the nearest neighbors (Figure 1, S5 – 15; Figure 3, 4, S4 – 15; Figure S1 – 30; Figure S3 – 20). BBKNN was implemented when necessary and calculated using the top 50 principal components with 3 neighbors per batch. The UMAP algorithm was then applied for visualization on 2 dimensions. Using the Louvain algorithm, clusters were identified at set resolutions (Figure 1, 3, 4, S3, S4, S5 – 0.4; Figure S1 – 0.25).

#### Cell Scoring

Cells were scored based on expression of a set of 50-60 marker genes per cell type. Gene lists were compiled based on the previously-published top 50-60 most differentially expressed genes from *in vivo* cells of the cell type of interest (Miller *et al*., 2020). See Table S1 for gene lists. After obtaining the log normalized and scaled expression values for the data set, scores for each cell were calculated as the average z-score within each set of selected genes.

#### Cluster Annotation

Using canonically expressed gene markers, each cluster’s general cell identity was annotated. The list of genes can be found in Figure S1A.

#### Sub-clustering

After annotating clusters within the UMAP embedding, specific clusters of interest were identified for further sub-clustering and analysis. The corresponding cells were extracted from the original filtered but unnormalized data matrix to include 35,561 cells in Figure 1, and S5, 2,312 cells for Figure 3A-C, and 749 cells for Figure 3D-G, 4, and S4. The extracted cell matrix then underwent log normalization, variable gene extraction, linear regression, z transformation, and dimension reduction to obtain a 2-dimensional UMAP embedding for visualization.

### Tissue Processing, Staining, and Quantification

All fluorescent images were taken using a NIKON A1 confocal microscope, an Olympus IX83 fluorescence microscope, or an Olympus IX71 fluorescence inverted microscope and were assembled using Photoshop CC 2021. Imaging parameters were kept consistent for images in the same experiment and post-image processing was performed equally on all images in the same experiment.

#### Tissue Processing

Tissue was immediately fixed in 10% Neutral Buffered Formalin for 24 hours at room temperature on a rocker, washed 3x, for 15 minutes each, with UltraPure DNase/RNase-Free Distilled Water (Thermo Fisher, Cat#10977015), and dehydrated for 1 hour in each of the following alcohol series diluted in UltraPure DNase/RNase-Free Distilled Water: 25% MeOH, 50% MeOH, 75% MeOH, 100% MeOH, 100% EtOH, 70% EtOH. Tissue was processed into paraffin blocks in an automated tissue processor (Leica ASP300) with 1-hour solution changes. For FISH, all equipment was sprayed with RNase AWAY (Thermo Fisher, Cat#700511) prior to sectioning. Paraffin blocks were sectioned into 4µm-thick sections for FISH (no longer than one week prior to performing FISH) or 4-7µm-thick sections for IF onto charged glass slides. Slides were baked for 1 hour in a 60°C dry oven (within 24 hours of performing FISH). Slides were stored at room temperature in a slide box containing a silicone desiccator packet and with the seams sealed with parafilm.

#### IF Protein Staining

Tissue slides were rehydrated in Histo-Clear II (National Diagnostics, Cat#HS-202) 2x for 5 minutes each, then put through the following solutions for 2x for 2 minutes each: 100% EtOH, 95% EtOH, 70% EtOH, 30% EtOH. Then, slides were put in double-distilled water (ddH20) 2x for 5 minutes each. Antigen retrieval was performed by steaming slides in 1X Sodium Citrate Buffer (100mM trisodium citrate (Sigma, Cat#S1804), 0.5% Tween 20 (Thermo Fisher, Cat#BP337), pH 6.0) for 20 minutes and subsequently cooling and washing quickly (moving slides up and down 5x) 2x in ddH20 and 2x in 1X PBS. Slides were incubated in a humidified chamber at room temperature for 1 hour with blocking solution (5% normal donkey serum (Sigma, Cat#D9663) in PBS with 0.1% Tween 20). Slides were then incubated in primary antibodies diluted in blocking solution in a humidified chamber at 4°C overnight. Slides were washed 3x in 1X PBS for 5 minutes each. Slides were incubated with secondary antibodies and DAPI (1µg/mL) diluted in blocking solution and placed in a humidified chamber at room temperature for 1 hour, then were washed 3x in 1X PBS for 5 minutes each. Slides were mounted in ProLong Gold (Thermo Fisher, Cat#P369300) and imaged within 2 weeks. Stained slides were stored in the dark a 4°C. All primary antibody concentrations are listed in Table S2. Secondary antibodies were raised in donkey, purchased from Jackson Immuno, and were used at a dilution of 1:500.

#### FISH

The FISH protocol was performed according to the manufacturer’s instructions (ACD Bio; RNAscope multiplex fluorescent manual protocol) with a 6-minute protease treatment and 15-minute antigen retrieval. For IF protein co-stains, the last step of the FISH protocol (DAPI) was skipped. Instead, the slides were washed 1x in PBS followed by the IF protocol above, beginning with the blocking step.

#### Quantification of IF and FISH images

All FISH images for quantification were taken at 40x magnification. For IF, 20x or 40x magnification was used. Nuclear stains and punctate FISH stains were analyzed using unbiased automated signal detection and quantification using CellProfiler (Lamprecht, Sabatini and Carpenter, 2007; Jones *et al*., 2008; Erben *et al*., 2018). Punctate per image, number of cells per image, punctate associated with specific nuclear stains, and numbers of specific positive nuclear stains were quantified using CellProfiler. Statistical analysis was performed using ordinary one-way ANOVA or Welch’s t test.

### RNA Extraction and qRT-PCR

Three biological replicates as well three technical replicates from the same biological specimen were included in each analysis. mRNA was isolated using the MagMAX-96 Total RNA Isolation Kit (Thermo Fisher, Cat#AM1830) or the PicoPure RNA Isolation Kit (Thermo Fisher, Cat#KIT0204) for FACS-sorted cells, and RNA quality and yield was measured on a Nanodrop 2000 spectrophometer just prior to cDNA synthesis. cDNA synthesis was performed using 100ng RNA from each sample and using the SuperScript VILO cDNA Kit (Thermo Fisher, Cat#11754250). qRT-PCR was performed on a Step One Plus Real-Time PCR System (Thermo Fisher, Cat#43765592R) using QuantiTect SYBR Green PCR Kit (Qiagen, Cat# 204145). Primer sequences can be found in Table S2. Expression of genes in the measurement of arbitrary units was calculated relative to GAPDH using the following equation:

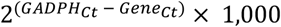

## Quantification and Statistical Analysis

Graphs and statistical analysis for RT-qPCR and FISH/IF quantification were performed in GraphPad Prism software. Quantification of FISH and IF were done using CellProfiler software. See figure legends for the number of replicates used, the statistical test performed, and the p-values used to determine significance (if p-values are not reported in the figure) for each analysis.

## Notes

https://github.com/jason-spence-lab/Hein_2021

### Competing Interest Statement

The authors have declared no competing interest.

