## Supplemental Figures and Tables for "R-SPONDIN2^+^ Mesenchymal Cells Form the Bud Tip Progenitor Niche During Human Lung Development"

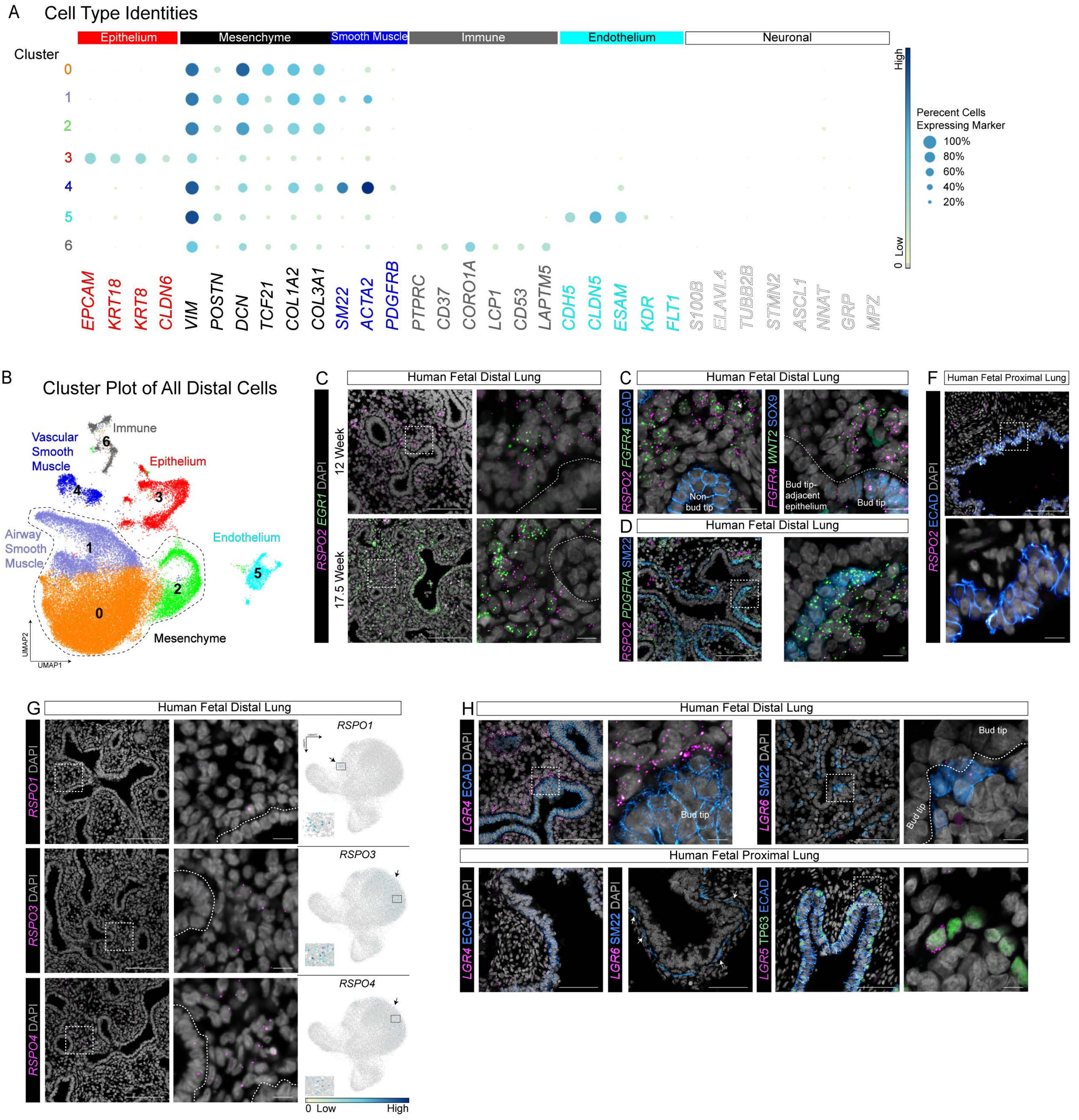

A Cell Type Identities
Epithelium
Mesenchyme
Smooth Muscle
Immune
Endothelium
Neuronal
Cluster
0
1
2
3
4
5
6
EPCAM KRT18 KRT8 CLDN6 VIM POSTN DCN TCF21 COL1A2 COL3A1 SM22 ACTA2 PDGFRB PTPRC CD37 CORO1A LCP1 CD53 LAPTM5 CDH5 CLDN5 ESAM KDR FLT1 S100B ELAVL4 TUBB2B STMN2 ASCL1 NNAT GRP MPZ
High
Low
0
Perecent Cells
Expressing Marker
100%
80%
60%
40%
20%
B Cluster Plot of All Distal Cells
Immune
Vascular Smooth Muscle
Epithelium
Airway Smooth Muscle
Endothelium
Mesenchyme
UMAP1
UMAP2
C Human Fetal Distal Lung
RSPO2 EGR1 DAPI
12 Week
17.5 Week
C Human Fetal Distal Lung
RSPO2 FGFR4 ECAD
Non-bud tip
FGFR4 WNT2 SOX9
Bud tip-adjacent epithelium
Bud tip
D Human Fetal Distal Lung
RSPO2 PDGFRA SM22
F Human Fetal Proximal Lung
RSPO2 ECAD DAPI
G Human Fetal Distal Lung
RSPO1 DAPI
RSPO1
RSPO3 DAPI
RSPO3
RSPO4 DAPI
RSPO4
0 Low High
H Human Fetal Distal Lung
LGR4 ECAD DAPI
Bud tip
LGR6 SM22 DAPI
Bud tip
Bud tip
Human Fetal Proximal Lung
LGR4 ECAD DAPI
LGR6 SM22 DAPI
LGR5 TP63 ECAD

**Supplementary Figure 1. Identification of major cell classes in the distal lung, characterization of the distal lung mesenchymal populations, and *RSPO* and *LGR* expression patterns**

(A) Dot plot of genes enriched in each cluster shown in Figure S1B. The dot size represents the percentage of cells expressing the gene in the corresponding cluster and the color indicates log-normalized and z-transformed expression level of the gene. Clusters are colored corresponding to the cluster plot in Figure S1B. Each cluster is associated with a major cell class (epithelium, mesenchyme, immune, endothelium, and neuronal).

(B) Cluster plot of cells from all time points sequenced using single cell RNA sequencing. Each dot represents a single cell and cells were computationally clustered based on transcriptional similarities. The plot is colored and numbered by cell-type identity of the cells composing each cluster. Cell-type labels for each cluster are based on expression of canonical cell-type markers displayed in the dot plot in Figure S1A. Cells from the epithelium, mesenchyme, endothelium, and immune system were identified.

(C) Multiplexed fluorescence *in situ* hybridization (FISH) of *RSPO2* and *EGR1*, a marker distinguishing two non-smooth muscle mesenchymal cell clusters (clusters 0 and 1) in Figure 1B, on 12- and 17.5-week human fetal distal lung tissue sections. Dotted lines outline epithelium. DAPI is shown in gray. Scale bars represent 100µm or 10µm for insets. Expression of *EGR1* is rare in the 12-week tissue but can be found in a small subset of *RSPO2*$^+$ mesenchymal cells and in some epithelial cells. By 17.5 weeks, *EGR1* expression is much stronger. Many *EGR1*$^+$/*RSPO2*$^+$ mesenchymal cells and *EGR1*$^-$/*RSPO2*$^+$ mesenchymal cells were found, but *EGR1*$^+$/*RSPO2*$^-$ mesenchymal cells were undetectable. *EGR1* expression was found throughout the epithelium, with expressing increasing from distal to proximal epithelium.

(D) Multiplexed FISH of *FGFR4* and *WNT2*, markers co-expressed in *RSPO2*$^+$ mesenchymal cells. The leftmost image shows a multiplexed FISH for *RSPO2* and *FGFR4* combined with immunofluorescence (IF) for the pan-epithelial marker ECAD on 13-week human fetal distal lung tissue sections. DAPI is shown in gray. Scale bars represent 100µm or 10µm for insets. The rightmost image shows a multiplexed FISH of *FGFR4* and *WNT2* combined with IF for the bud tip marker SOX9 on 12-week human fetal distal lung tissue sections. DAPI is shown in gray. Scale bars represent 10µm. *RSPO2*, *FGFR4*, and *WNT2* are co-expressed in the mesenchyme. *FGFR4* is also expressed in bud tip progenitor cells.

(E) Multiplexed FISH of *PDGFRA* and *RSPO2* and co-IF for SM22 on 12-week human fetal distal lung tissue sections. DAPI is shown in gray. Scale bars represent 100µm or 10µm for insets. *PDGFRA* is expressed broadly through most of the distal lung mesenchyme and appears enriched in SM22$^+$ cells directly adjacent to bud tip regions.

(F) FISH of *RSPO2* and co-IF for the pan-epithelial marker ECAD on 17.5-week human distal proximal lung tissue sections. DAPI is shown in gray. Scale bars represent 100µm or 10µm for insets. *RSPO2* is largely absent from both proximal epithelium and proximal mesenchyme.

(G) FISH and UMAP feature plots of *RSPO1* (top), *RSPO3* (middle), and *RSPO4* (bottom) on 15-week human fetal distal lung tissue sections. For FISH, DAPI is shown in gray and scale bars represent 100µm or 10µm for insets, and dotted lines outline epithelium. The UMAP feature plots correspond to the cluster plot in Figure 1B, and the color of each dot indicates log-normalized and z-transformed expression level for the marker listed in the represented cell. In general, *RSPO1*, *RSPO3*, and *RSPO4* are expressed in the same mesenchymal cell cluster(s) as *RSPO2* in the distal lung but are expressed in fewer cells with no clear expression pattern. They appear to have low to no expression in the epithelium.

(H) FISH of *LGR5* in the proximal lung and *LGR4* and *LGR6* in the distal and proximal lung and co-IF for the pan-epithelial marker ECAD, smooth muscle marker SM22, and/or the basal cell marker TP63 on 12-17-week human fetal distal lung tissue sections. DAPI is shown in gray. Scale bars represent 100µm or 10µm for insets. *LGR4* and *LGR6* are not expressed in bud tip progenitor cells. *LGR4* is expressed broadly throughout the proximal and distal mesenchyme and *LGR6* expression is localized to airway smooth muscle cells. *LGR5* is expressed in a subset of basal cells in the proximal lung.

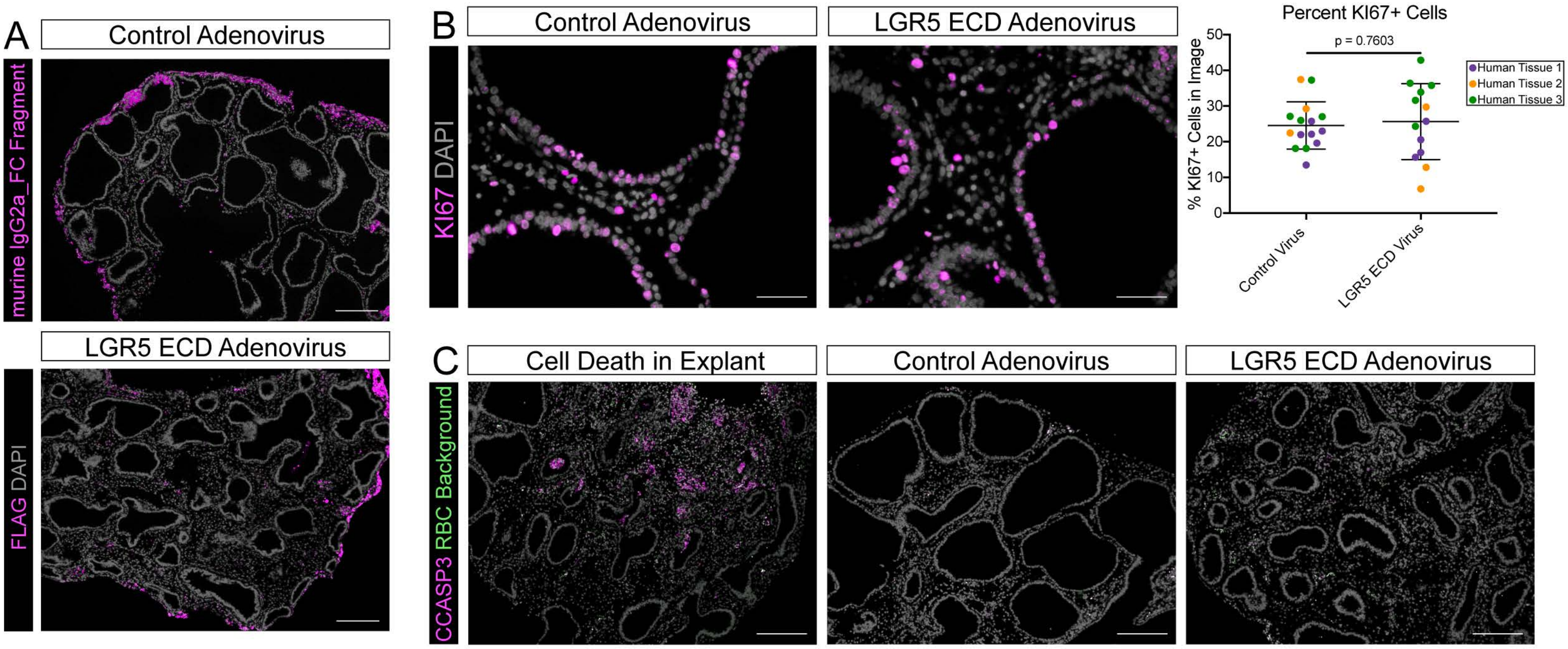
A
Control Adenovirus
murine IgG2a_FC Fragment
LGR5 ECD Adenovirus
FLAG DAPI
B
Control Adenovirus
LGR5 ECD Adenovirus
KI67 DAPI
Percent KI67+ Cells
p = 0.7603
% KI67+ Cells in Image
0
10
20
30
40
50
Control Virus
LGR5 ECD Virus
Human Tissue 1
Human Tissue 2
Human Tissue 3
C
Cell Death in Explant
Control Adenovirus
LGR5 ECD Adenovirus
CCASP3 RBC Background

**Supplementary Figure 2. Confirmation of adenovirus infection and proliferation versus apoptosis in lung explants**

(A) Immunofluorescence (IF) staining for murine IgG2a_FC on the control adenovirus (ad)-infected explants detects the presence of the control ad in explants and IF staining for FLAG on the LGR5 ECD ad-infected explants detects the presence of the LGR5 ECD ad in explants. DAPI is shown in gray. Scale bars represent 200μm.

(B) KI67 immunofluorescence staining on and quantification of sections from control adenovirus (ad)-infected explants and LGR5 ECD ad-infected explants. DAPI is shown in gray. Scale bars represent 50μm. The number of proliferative cells, measured by KI67 staining, is not significantly different between control ad-infected explants and LGR5 ECD ad-infected explants, with 24.6% of cells staining positive for KI67 in control ad-infected explants and 25.6% of cells staining positive for KI67 in LGR5 ECD ad-infected explants ($p = 0.76030$, Welch's t test). This quantification was performed in three unique biological samples with one to three technical replicates and a minimum of three image fields for each sample.

(C) CCASP3 immunofluorescence staining on sections from an explant that grew abnormally large as well as typical-sized control and LGR5 ECD ad-infected explants. DAPI is shown in gray. Scale bars represent 200μm. Large explants often had visible cell death near their center. Cell death in average-sized control and LGR5 ECD ad-infected explants was comparably low. Background from red blood cells (RBC) is shown in green in order to identify true CCASP3 staining.

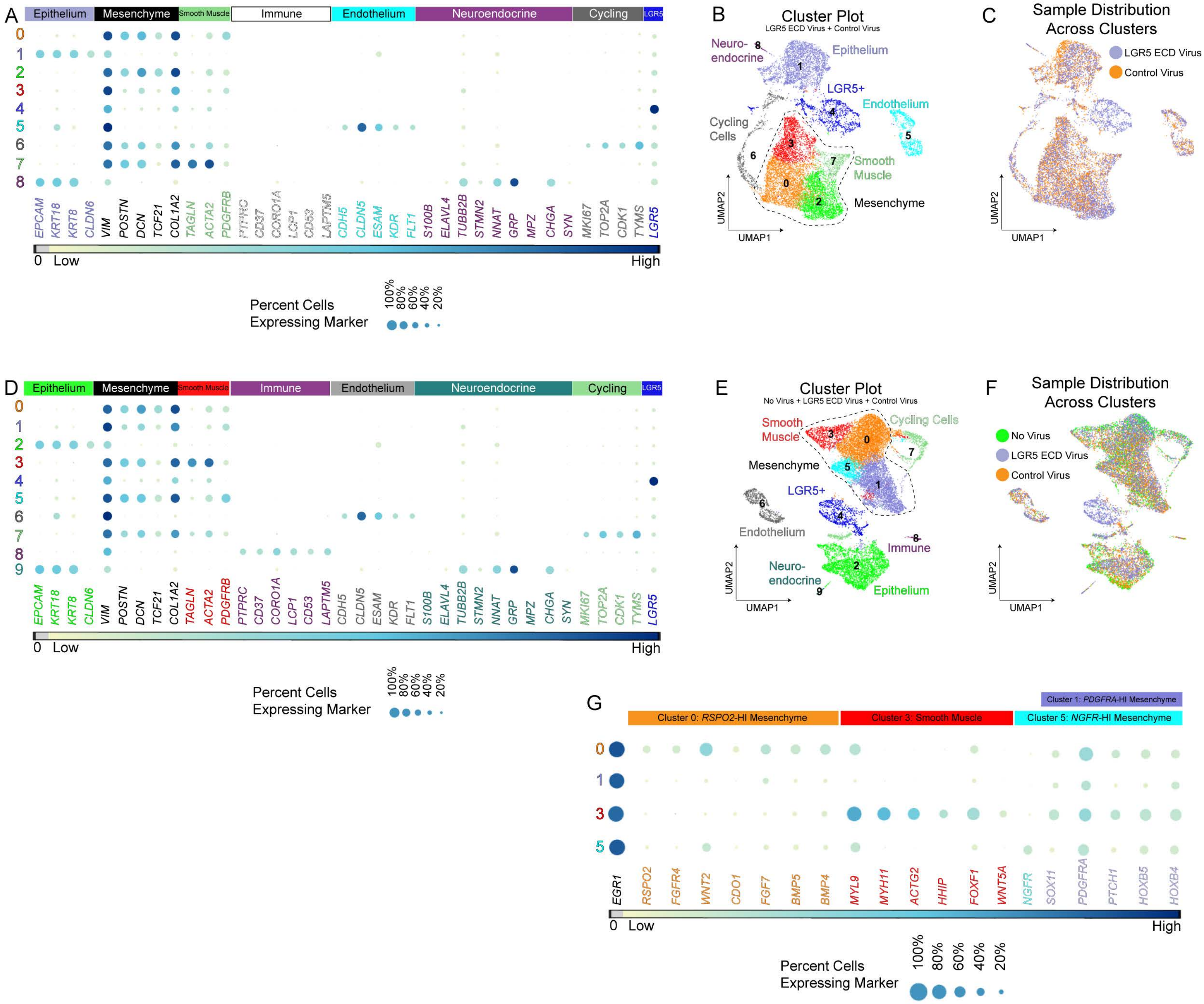
A
Epithelium
Mesenchyme
Smooth Muscle
Immune
Endothelium
Neuroendocrine
Cycling
LGR5
EPCAM KRT18 KRT8 CLDN6 VIM POSTN DCN TCF21 COL1A2 TAGLN ACTA2 PDGFRB PTPRC CD37 CORO1A LCP1 CD53 LAPTM5 CDH5 CLDN5 ESAM KDR FLT1 S100B ELAVL4 TUBB2B STMN2 NNAT GRP MPZ CHGA SYN MKI67 TOP2A CDK1 TYMS LGR5
0 Low
High
Percent Cells
Expressing Marker
100%
80%
60%
40%
20%
B
Cluster Plot
LGR5 ECD Virus + Control Virus
Neuro-
endocrine
Epithelium
LGR5+
Endothelium
Cycling
Cells
Smooth
Muscle
Mesenchyme
UMAP2
UMAP1
C
Sample Distribution
Across Clusters
LGR5 ECD Virus
Control Virus
UMAP2
UMAP1
D
Epithelium
Mesenchyme
Smooth Muscle
Immune
Endothelium
Neuroendocrine
Cycling
LGR5
EPCAM KRT18 KRT8 CLDN6 VIM POSTN DCN TCF21 COL1A2 TAGLN ACTA2 PDGFRB PTPRC CD37 CORO1A LCP1 CD53 LAPTM5 CDH5 CLDN5 ESAM KDR FLT1 S100B ELAVL4 TUBB2B STMN2 NNAT GRP MPZ CHGA SYN MKI67 TOP2A CDK1 TYMS LGR5
0 Low
High
Percent Cells
Expressing Marker
100%
80%
60%
40%
20%
E
Cluster Plot
No Virus + LGR5 ECD Virus + Control Virus
Smooth
Muscle
Cycling Cells
Mesenchyme
LGR5+
Endothelium
Immune
Neuro-
endocrine
Epithelium
UMAP2
UMAP1
F
Sample Distribution
Across Clusters
No Virus
LGR5 ECD Virus
Control Virus
UMAP2
UMAP1
G
Cluster 1: PDGFRA-HI Mesenchyme
Cluster 0: RSPO2-HI Mesenchyme
Cluster 3: Smooth Muscle
Cluster 5: NGFR-HI Mesenchyme
EGR1 RSPO2 FGFR4 WNT2 CDO1 FGF7 BMP5 BMP4 MYL9 MYH11 ACTG2 HHIP FOXF1 WNT5A NGFR SOX11 PDGFRA PTCH1 HOXB5 HOXB4
0 Low
High
Percent Cells
Expressing Marker
100%
80%
60%
40%
20%

**Supplementary Figure 3. Identification of major cell classes in lung explants infected with LGR5 ECD or control adenovirus and non-infected explants**

A) Dot plot of genes enriched in each cluster shown in Figure S3B. The dot size represents the percentage of cells expressing the gene in the corresponding cluster, and the dot color indicates log-normalized and z-transformed expression level of the gene. Clusters are colored corresponding to the cluster plot in Figure S3B.

B) Cluster plot of cells from LGR5 ECD adenovirus (ad)-infected explants and control ad-infected explants sequenced using single cell RNA sequencing. Each dot represents a single cell and cells were computationally clustered based on transcriptional similarities. The plot is colored and numbered by cell-type identity of the cells composing each cluster. Cell-type labels for each cluster are based on expression of canonical cell-type markers displayed in the dot plot in Figure S3A. Cells from the epithelium, mesenchyme, and endothelium were identified. Also identified was a cluster of neuro-endocrine cells (positive expression for epithelial markers, neuronal markers, and neuro-endocrine markers *CHGA* and *SYN*), a cluster of cycling cells, and a cluster of cells defined by high expression of *LGR5*.

C) Cluster plot corresponding to Figure S3B. Each dot represents a single cell and dots/cells are colored by the sample from which they came from.

D) Dot plot of genes enriched in each cluster shown in Figure S3E. The dot size represents the percentage of cells expressing the gene in the corresponding cluster, and the dot color indicates log-normalized and z-transformed expression level of the gene. Clusters are colored corresponding to the UMAP plot in Figure S3E.

E) Cluster plot of cells from non-viral infected explants, LGR5 ECD ad-infected explants, and control ad-infected explants sequenced using single cell RNA sequencing. Each dot represents a single cell and cells were computationally clustered based on transcriptional similarities. The plot is colored and numbered by cell-type identity of the cells composing each cluster. Cell-type labels for each cluster are based on expression of canonical cell-type markers displayed in the dot plot in Figure S3D. Cells from the epithelium, mesenchyme, endothelium, and immune system were identified. Also identified was a cluster of neuro-endocrine cells (positive expression for epithelial markers, neuronal markers, and neuro-endocrine markers *CHGA* and *SYN*), a cluster of cycling cells, and a cluster of cells defined by high expression of *LGR5*.

F) UMAP plot corresponding to the cluster plot in Figure S3E. Each dot represents a single cell and dots/cells are colored by the sample from which they came from.

G) Dot plot of mesenchymal cell type markers that were identified in Figure 1D. Clusters are colored corresponding to the cluster plot in Figure S3E. The dot size represents the percentage of cells expressing the marker in the corresponding cluster, and the dot color indicates log-normalized and z-transformed expression level of the marker. The explants retained the general mesenchymal cell populations found in the fetal lung with small changes in expression of some genes in the non-smooth muscle mesenchymal cell populations. Expression of *EGR1* increased in every mesenchymal cell cluster compared to the *in vivo* fetal lung.

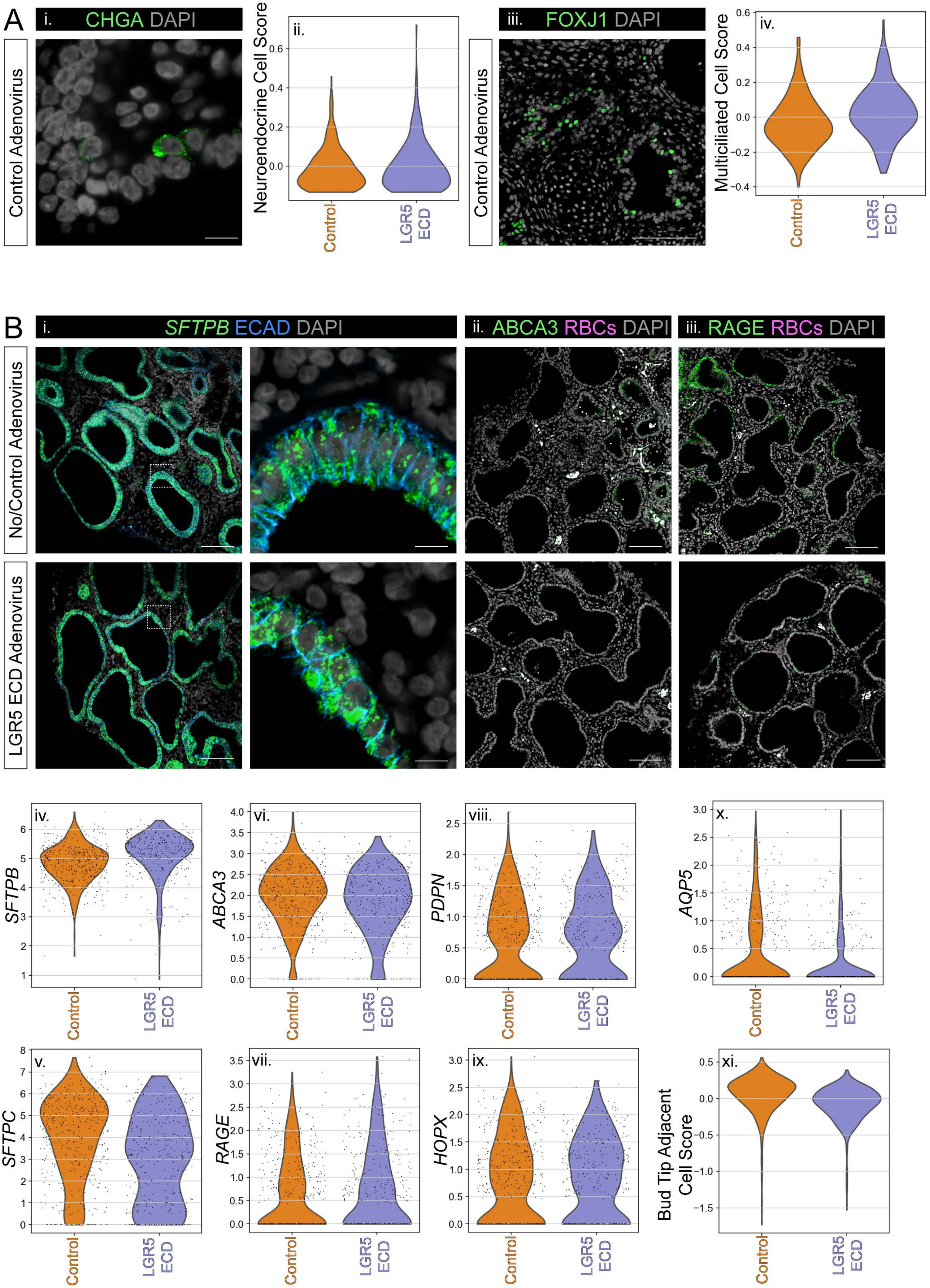
A
i. CHGA DAPI
Control Adenovirus
ii.
Neuroendocrine Cell Score
Control
LGR5 ECD
iii. FOXJ1 DAPI
Control Adenovirus
iv.
Multiciliated Cell Score
Control
LGR5 ECD
B
i. SFTPB ECAD DAPI
ii. ABCA3 RBCs DAPI
iii. RAGE RBCs DAPI
No/Control Adenovirus
LGR5 ECD Adenovirus
iv. SFTPB
v. SFTPC
vi. ABCA3
vii. RAGE
viii. PDPN
ix. HOPX
x. AQP5
xi. Bud Tip Adjacent Cell Score
Control
LGR5 ECD

**Supplementary Figure 4. Inhibition of RSPO2-potentiated WNT signaling in lung explants results in minimal bud tip differentiation into proximal neuroendocrine and multicilated cell types and distal cell types**

(A) Expression of proximal neuroendocrine and multiciliated cell type markers in LGR5 ECD ad-infected explants and control ad-infected explants. For immunofluorescence (IF) images, DAPI is shown in gray. Scale bars represent 10μm for CHGA and 100μm for FOXJ1. For violin plots, cell score for the listed proximal cell type was calculated from single cell RNA sequencing (scRNA-seq) data as the average expression of the top 50 enriched genes in *in vivo* cells of the listed cell type (see methods). **(i.)** Immunofluorescence (IF) staining for the neuroendocrine cell marker CHGA on a section from a control ad-infected explant. $CHGA^+$ cells were rare in both explant conditions. **(ii.)** Cell score for neuroendocrine cells is similar between the two conditions, with a small increase in the LGR5 ECD ad-infected group. **(iii.)** IF staining for the multiciliated cell marker FOXJ1 on a section from a control ad-infected explant. $FOXJ1^+$ cells were only found in proximal airway structures in explants from both conditions. **(iv.)**. Cell score for multiciliated cells is higher in cells from the LGR5 ECD ad-infected explants compared to cells from the control ad-infected explants.

(B) Expression of distal (alveolar/bud tip adjacent) cell type markers in LGR5 ECD ad-infected explants and control ad-infected explants or non-viral infected explants. Non-viral infected tissue was used for antibodies raised in mice to avoid secondary antibodies from detecting the control adenovirus and misleading interpretation of the stain. For IF and fluorescence *in situ* hybridization (FISH), DAPI is shown in gray. Scale bars represent 100μm or 10μm for insets. **(i.)** FISH for *SFTPB*, expressed in distal cell types as well as some proximal cell types, on sections from control ad-infected explants and LGR5 ECD ad-infected explants. *SFTPB* mRNA staining is high in all epithelium from both conditions. **(ii. – iii.)** IF staining for the alveolar type II and type I markers ABCA3 and RAGE, respectively, on sections from non-infected explants and LGR5 ECD ad-infected explants. The pink channel shows autofluorescence from red blood cells (RBSc) that overlap with autofluorescence in the green channel in order to more clearly visualize true antibody staining. At the protein level, both of these markers appear moderately increased in non-infected explants compared to LGR5 ECD ad-infected explants. **(iv. – x.)** Violin plots displaying log-normalized and z-transformed expression level of a single gene from scRNA-seq on control ad-infected explants and LGR5 ECD ad-infected explants. At the mRNA level, expression of *SFTPB* and *RAGE* is moderately increased in LGR5 ECD ad-infected, expression of *SFTPC* and *AQP5* is moderately increased in control ad-infected explants, and expression of *ABCA3*, *PDPN*, and *HOPX* is similar between the two conditions. **(xi.)** Violin plot displaying bud tip adjacent cell score, calculated from scRNA-seq data as the average expression of the top 50 genes enriched in *in vivo* bud tip adjacent cells (see methods). Cell score for bud tip adjacent cells is higher in cells from the control ad-infected explants compared to cells from the LGR5 ECD ad-infected explants.

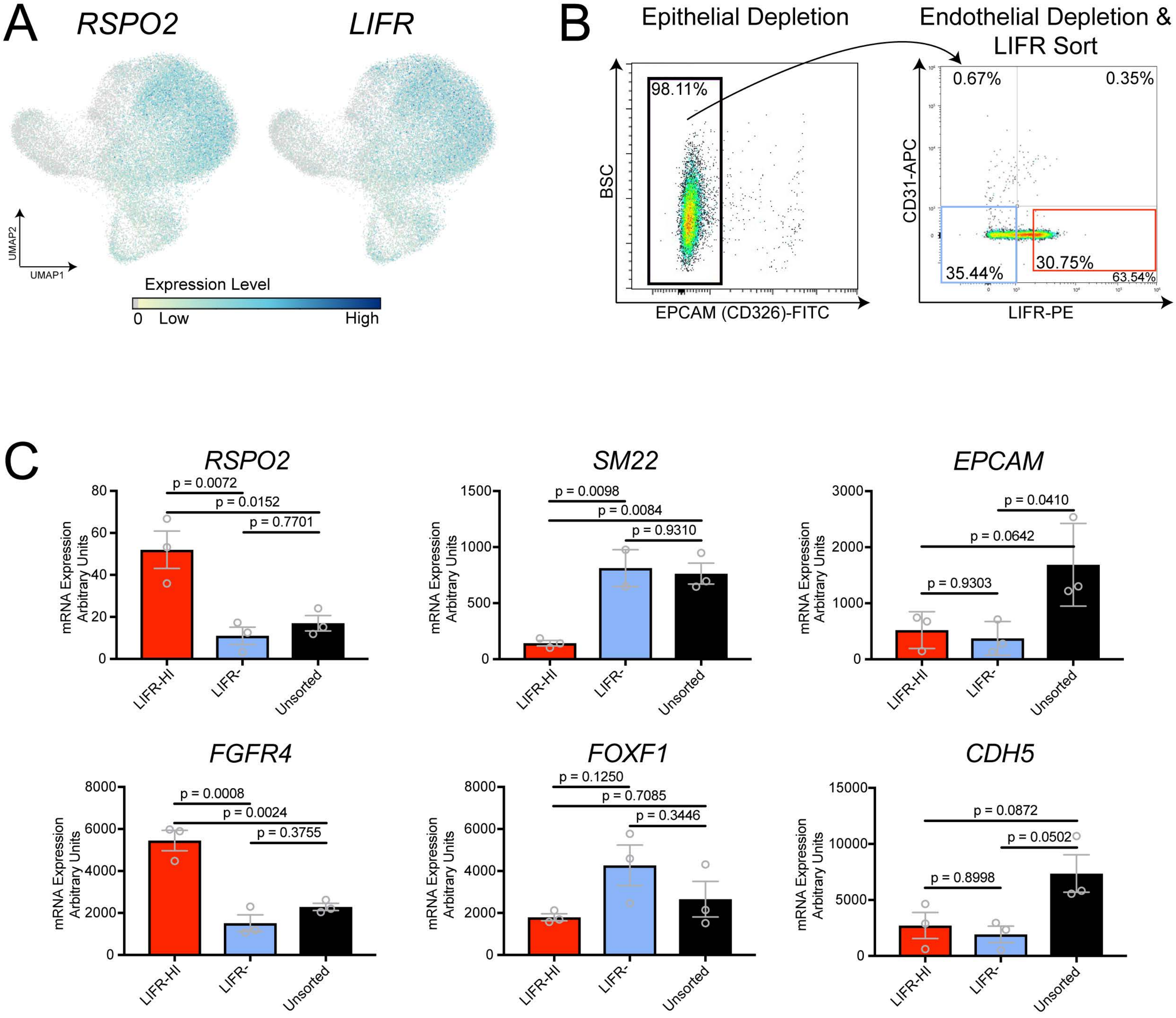
A
RSPO2
LIFR
UMAP2
UMAP1
Expression Level
0 Low
High
B
Epithelial Depletion
Endothelial Depletion &
LIFR Sort
98.11%
BSC
EPCAM (CD326)-FITC
0.67%
0.35%
CD31-APC
35.44%
30.75%
63.54%
LIFR-PE
C
RSPO2
p = 0.0072
p = 0.0152
p = 0.7701
mRNA Expression Arbitrary Units
LIFR-HI
LIFR-
Unsorted
SM22
p = 0.0098
p = 0.0084
p = 0.9310
EPCAM
p = 0.0410
p = 0.0642
p = 0.9303
FGFR4
p = 0.0008
p = 0.0024
p = 0.3755
FOXF1
p = 0.1250
p = 0.7085
p = 0.3446
CDH5
p = 0.0872
p = 0.0502
p = 0.8998

**Supplementary Figure 5. FACS of RSPO2⁺ and SM22⁺ mesenchymal cell populations**

A) UMAP feature plots corresponding to the cluster plot in Figure 1B and displaying expression levels *RSOP2* and *LIFR*. Each dot represents a single cell, and the color of each dot indicates log-normalized and z-transformed expression level of the given gene in the represented cell. *RSPO2* and *LIFR* appear strongly co-expressed in the mesenchyme.

B) Flow cytometric analysis and pipeline for fluorescence activated cell sorting (FACS) of $RSPO2^+$ and $SM22^+$ mesenchymal cell populations by LIFR. Representative flow cytometry plots for EPCAM (CD326) (left plot) as well as CD31 and LIFR (right plot) from one tissue sample. After gating of live, single cells, epithelial cells were negatively selected for via EPCAM. After negative selection of $EPCAM^+$ cells, endothelial cells were separated from mesenchyme via CD31. The $CD326^-$/$CD31^-$/$LIFR^{HI}$ cells (Q4, red box) and $CD326^-$/$CD31^-$/$LIFR^-$ cells (Q3, blue box) were collected for co-culture experiments. $LIFR^{HI}$ cells were defined as the top 30% of cells expressing LIFR. Isotype controls were used to set quadrants, and individual antibody/color channel controls were used to set compensation.

C) qPCR from cells sorted using the FACS pipeline shown in Figure S5B. Genes expected to be expressed uniquely in *RSPO2*$^+$ mesenchymal cells (*RSPO2*, *FGFR4*) were enriched in the $CD326^-$/$CD31^-$/$LIFR^{HI}$ population while genes expected to be expressed uniquely in airway smooth muscle cells (*SM22*, *FOXF1*) were enriched in the $CD326^-$/$CD31^-$/$LIFR^-$ population. Epithelial cells (*EPCAM*$^+$) and endothelial cells (*CDH5*$^+$) cells were successfully depleted from both sorted populations. p-values were calculated using a one-way ANOVA followed by Turkey's multiple comparison test to compare the mean of each group with the mean of every other group. Error bars show standard deviation. Each data point represents a unique biological replicate from three separate experiments.

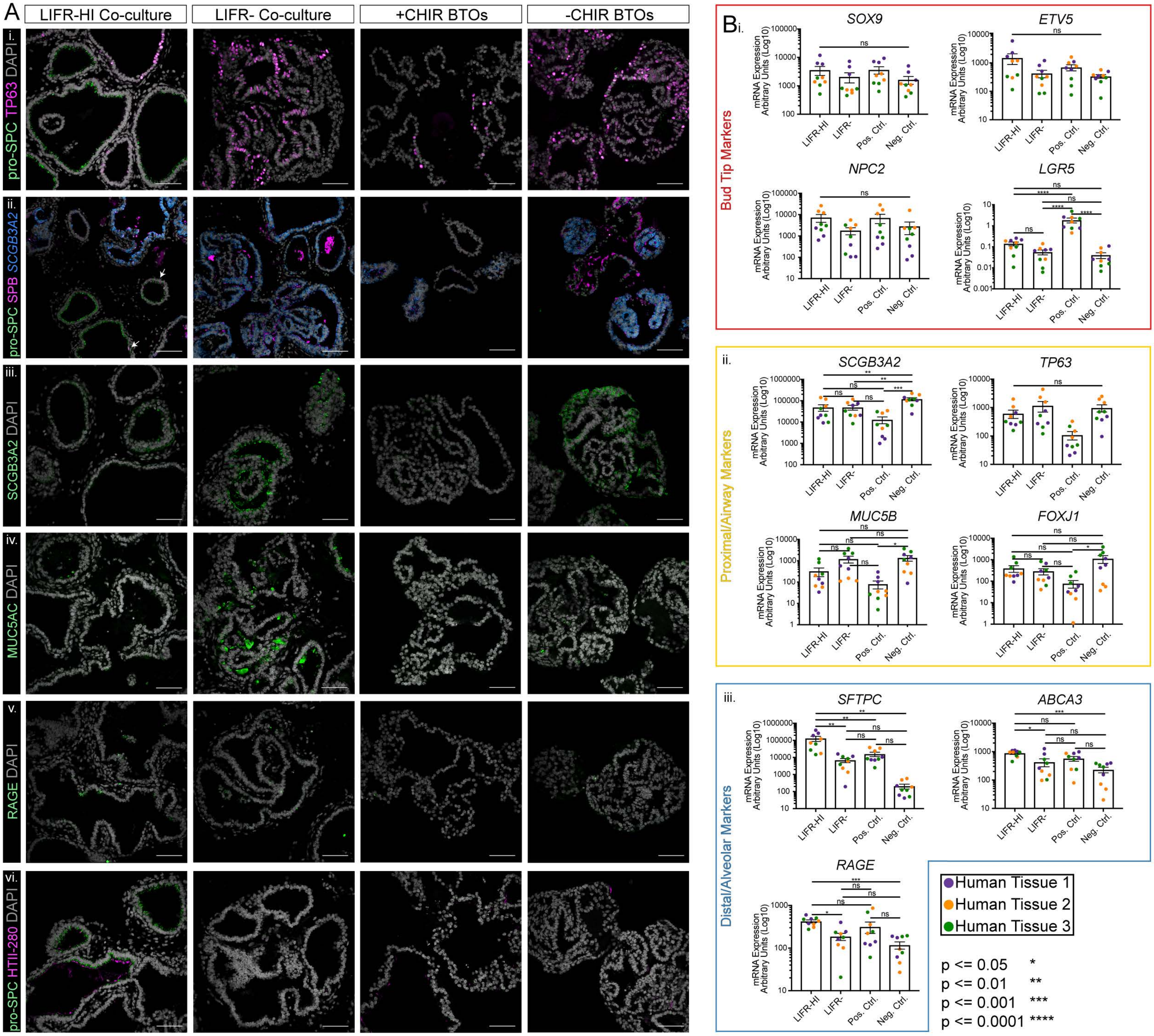
A
LIFR-HI Co-culture
LIFR- Co-culture
+CHIR BTOs
-CHIR BTOs
i. pro-SPC TP63 DAPI
ii. pro-SPC SPB SCGB3A2
iii. SCGB3A2 DAPI
iv. MUC5AC DAPI
v. RAGE DAPI
vi. pro-SPC HTII-280 DAPI
B i.
Bud Tip Markers
SOX9
ETV5
NPC2
LGR5
ii.
Proximal/Airway Markers
SCGB3A2
TP63
MUC5B
FOXJ1
iii.
Distal/Alveolar Markers
SFTPC
ABCA3
RAGE
mRNA Expression Arbitrary Units (Log10)
LIFR-HI
LIFR-
Pos. Ctrl.
Neg. Ctrl.
Human Tissue 1
Human Tissue 2
Human Tissue 3
p <= 0.05 *
p <= 0.01 **
p <= 0.001 ***
p <= 0.0001 ****

**Supplementary Figure 6. Differentiated proximal and distal cell type markers in bud tip & mesenchyme co-cultures**

**(A)** Expression of proximal and distal differentiated cell type markers on tissue sections from $LIFR^{HI}$ co-cultures, $LIFR^{-}$ co-cultures, positive control (+CHIR99021) bud tips, and negative control (-CHIR99021) bud tips after 11 days of culture from three independent experiments. Scale bars represent 100µm. **(i.)** Immunofluorescence (IF) for distal/alveolar marker pro-SFTPC and basal cell marker TP63. Pro-$SFTPC^{+}$ cells were only detected in $LIFR^{HI}$ co-cultures. The positive control bud tips were the only condition with few $TP63^{+}$ cells. **(ii.)** Fluorescence *in situ* hybridization for secretory cell marker *SCGB3A2* with co-IF for distal/alveolar markers pro-SFTPC and SFTPB. *SCGB3A2*$^{+}$ cells were detected in every condition. In the $LIFR^{-}$ co-culture and negative control bud tips, *SCGB3A2*$^{+}$ cells co-expressed high levels of SFTPB while in the $LIFR^{HI}$ co-culture and positive control bud tips, *SCGB3A2*$^{+}$ cells co-expressed low levels of SFTPB. Some pro-$SFTPC^{+}$ cells in the $LIFR^{HI}$ co-culture also expressed SFTPB. **(iii.)** IF for secretory cell marker SCGB3A2. At the protein level, SCGB3A2 was detected at high levels in $LIFR^{-}$ co-cultures and negative control bud tips, at low levels in the $LIFR^{HI}$ co-cultures, and was almost undetectable in positive control bud tips. **(iv.)** IF for goblet cell marker MUC5AC. At the protein level, MUC5AC was detected at high levels in $LIFR^{-}$ co-cultures, at low levels in the $LIFR^{HI}$ co-cultures and negative control bud tips, and was undetectable in positive control bud tips. **(v.)** IF for alveolar type I cell marker RAGE. More $RAGE^{+}$ cells were detected in $LIFR^{HI}$ co-culture compared to the other conditions. **(vi.)** IF for alveolar type II markers pro-SFTPC and HTII-280. Some pro-$SFTPC^{+}$ cells in the $LIFR^{HI}$ co-culture also expressed HTII-280, both of which were low or undetectable in all other conditions.

**(B)** RT-qPCR for bud tip, proximal/airway, and distal/alveolar markers on $LIFR^{HI}$ co-cultures, $LIFR^{-}$ co-cultures, positive control (+CHIR99021) bud tips, and negative control (-CHIR99021) bud tips after 11 days of culture from three independent experiments. Each color represents an independent experiment using bud tips and mesenchyme from unique specimens. Each data point of the same color represents a technical replicate from the same set of tissue specimens. Error bars represent standard error of the mean. Statistical tests were performed by ordinary one-way ANOVA followed by Dunnett's multiple comparison test. **(i.)** Bud tip genes *SOX9*, *ETV5*, and *NPC2* are highest in the $LIFR^{HI}$ co-culture and positive control bud tips, but not by a statistically significant amount. *LGR5* is statistically significantly higher in positive control bud tips compared to all other conditions. **(ii.)** Proximal airway genes *SCGB3A2* (secretory cells), *TP63* (basal cells), *MUC5B* (goblet cells), and *FOXJ1* (ciliated cells) are high in all conditions except for positive control bud tips. **(iii.)** Distal alveolar genes *SFTPC*, *ABCA3*, and *RAGE* are enriched in the $LIFR^{HI}$ condition compared to all other conditions.

**Supplementary Table 1. Gene lists for cell scoring analysis**
Genes used to calculate cell type scores in Figures 3, 4, and S4. The listed genes are the previously published top 50-56 differentially expressed genes in *in vivo* cells from the human fetal lung of the listed cell type on the tab (Miller *et al.*, 2020).

**Supplementary Table 2. Antibody and TSA dilutions and primer sequences**

| Antibody | Dilution |
|---|---|
| TAGLN (SM22) (Abcam) | 1:500 |
| SOX9 (R&D Systems)<br>(used in combination with FISH) | 1:500 |
| SOX9 (Millipore)<br>(used for IF only) | 1:500 |
| SOX9 (Santa Cruz)<br>(used for IF only) | 1:150 |
| ECAD (BD Biosciences) | 1:500 |
| SOX2 (R&D Systems) | 1:500 |
| TP63 (R&D Systems) | 1:500 |
| CHGA (Abcam) | 1:300 |
| RAGE (Abcam) | 1:500 |
| ABCA3 (Seven Hills Bioreagents) | 1:500 |
| FOXJ1 (Seven Hills Bioreagents) | 1:250 |
| FLAG (Sigma) | 1:250 |
| KI67 (Thermo Fisher) | 1:250 |
| CCAS3 (Cell Signaling) | 1:500 |
| Pro-SP-C (Seven Hills Bioreagents) | 1:500 |

| | |
|---|---|
| SP-B (Seven Hills Bioreagents) | 1:500 |
| SCGB3A2 (Abcam) | 1:500 |
| HTII-280 (Terrace Biotech) | 1:100 |
| Alpha smooth muscle actin (Sigma) | 1:500, 1 hour |
| MUC5AC (Abcam) | 1:500 |
| LIFR alpha PE (R&D Systems) | 10μL per $10^6$ cells |
| IgG1 PE isotype control (R&D Systems) | 10μL per $10^6$ cells |
| CD326 (EPCAM) FITC (Miltenyi) | 1:50 per $10^6$ cells |
| REA control IgG1 FITC (Miltenyi) | 1:50 per $10^6$ cells |
| CD31 APC (Miltenyi) | 1:50 per $10^6$ cells |
| REA control IgG1 APC (Miltenyi) | 0.6μL in 100μL for $10^6$ cells |
| CD31 647 (BD Biosciences) | 1:200 per $10^6$ cells |
| IgG2a FC control APC (Thermo Fisher) | 1:500 per $10^6$ cells |
| **Probe** | **TSA Plus Fluorophore & Dilution** |
| Hs-RSPO2-02 | TSA Plus Cyanine 3 (1:2500) |
| Hs-LGR5 | TSA Plus Cyanine 3 (1:2500) |
| Hs-AXIN2 | TSA Plus Cyanine 3 (1:2500) |
| Hs-PDGFRA-C2 | TSA Plus Cyanine 5 (1:2000) |

| | |
|---|---|
| Hs-FGFR4-no-XMm-C2 | TSA Plus Cyanine 5 (1:2000) |
| Hs-WNT2 | TSA Plus Cyanine 3 (1:2500) |
| Hs-EGR1-C2 | TSA Plus Cyanine 5 (1:2500) |
| Hs-RSPO1 | TSA Plus Cyanine 3 (1:2500) |
| Hs-RSPO3-01 | TSA Plus Cyanine 3 (1:2500) |
| Hs-RSPO4 | TSA Plus Cyanine 3 (1:2500) |
| Hs-LGR4 | TSA Plus Cyanine 3 (1:2500) |
| Hs-LGR6 | TSA Plus Cyanine 3 (1:2500) |
| Hs-SCGB3A2 | TSA Plus Cyanine 3 (1:2500) |
| Hs-SFTPB | TSA Plus Cyanine 3 (1:2500) |
| **Primer** | **Sequence** |
| SOX9 | F: GTACCCGCACTTGCACAAC<br>R: GTGGTCCTTCTTGTGCTGC |
| ETV5 | F: GAAGAGGTTCGCAGGGATAAG<br>R: TAAGGCCCAATCTACAGGTTTAC |
| LGR5 | F: CCTCAGCGTCTTCACCTCCT<br>R: TTTCTTTCCCAGGGAGTGGAT |
| NPC2 | F: CGGTTCTGTGGATGGAGTTAT<br>R: GGTGACATTGACGCTGTAAGA |
| SCGB3A2 | F: GGGGCTAAGGAAGTGTGTAAATG<br>R: CACCAAGTGTGATAGCGCCTC |
| TP63 | F: CCACAGTACACGAACCTGGG<br>R: CCGTTCTGAATCTGCTGGTCC |
| MUC5B | F: GTACCTGGACTGCAGCAACA<br>R: CTTGTAGGTGGCCTCGTTGT |

| | |
|---|---|
| FOXJ1 | F: CAACTTCTGCTACTTCCGCC<br>R: CGAGGCACTTTGATGAAGC |
| SFTPC | F: AGCAAAGAGGTCCTGATGGA<br>R: CGATAAGAAGGCGTTTCAGG |
| ABCA3 | F: TGCAGCGCCTACTTGAACTT<br>R: CTGAGCACAGCCATCGTCT |
| RAGE | F: TCCGGGCTGTGATGTTTTGA<br>R: CTCACCCACCCTGTCCAAAT |
| RSPO2 | F: TGAATGCCAGATGGTTTTGC<br>R: ATCTGCCGTGTTCTGGTTTC |
| FGFR4 | F: GAACCGCATTGGAGGCATT<br>R: TTCTCTACCAGGCAGGTGTATGTG |
| SM22 | F: AACAGCCTGTACCCTGATGG<br>R: CGGTAGTGCCCATCATTCTT |
| FOXF1 | F: AGTCCCCAATGCAAAGACAC<br>R: TCAGCAGAATTCCTGTGTGG |
| EPCAM | F: CTGCCAAATGTTTGGTGATG<br>R: CTTCTGACCCCAGCAGTGTT |
| CDH5 | F: CTTCACCCAGACCAAGTACACA<br>R: AATGGTGAAAGCGTCCTGGT |
| AXIN2 | F: AGTGTGAGGTCCACGGAAAC<br>R: CTGGTGCAAAGACATAGCCA |
| KI67 | F: TGGGTCTGTTATTGATGAGCC<br>R: TGACTTCCTTCCATTCTGAAGAC |
